# Chromatin remodeler BRG1 recruits huntingtin to repair DNA double-strand breaks in neurons

**DOI:** 10.1101/2024.09.19.613927

**Authors:** Subrata Pradhan, Keegan Bush, Nan Zhang, Raj K. Pandita, Chi-Lin Tsai, Charlene Smith, Devon F. Pandlebury, Sagar Gaikwad, Francis Leonard, Linghui Nie, Annie Tao, William Russell, Subo Yuan, Sanjeev Choudhary, Kenneth S. Ramos, Cornelis Elferink, Yogesh P. Wairkar, John A. Tainer, Leslie M. Thompson, Tej K. Pandita, Partha S. Sarkar

## Abstract

Persistent DNA double-strand breaks (DSBs) are enigmatically implicated in neurodegenerative diseases including Huntington’s disease (HD), the inherited late-onset disorder caused by CAG repeat elongations in Huntingtin (HTT). Here we combine biochemistry, computation and molecular cell biology to unveil a mechanism whereby HTT coordinates a Transcription-Coupled Non-Homologous End-Joining (TC-NHEJ) complex. HTT joins TC-NHEJ proteins PNKP, Ku70/80, and XRCC4 with chromatin remodeler Brahma-related Gene 1 (BRG1) to resolve transcription-associated DSBs in brain. HTT recruitment to DSBs in transcriptionally active gene-rich regions is BRG1-dependent while efficient TC-NHEJ protein recruitment is HTT-dependent. Notably, mHTT compromises TC-NHEJ interactions and repair activity, promoting DSB accumulation in HD tissues. Importantly, HTT or PNKP overexpression restores TC-NHEJ in a *Drosophila* HD model dramatically improving genome integrity, motor defects, and lifespan. Collective results uncover HTT stimulation of DSB repair by organizing a TC-NHEJ complex that is impaired by mHTT thereby implicating dysregulation of transcription-coupled DSB repair in mHTT pathophysiology.

**Highlights:**

• BRG1 recruits HTT and NHEJ components to transcriptionally active DSBs.
• HTT joins BRG1 and PNKP to efficiently repair transcription related DSBs in brain.
• Mutant HTT impairs the functional integrity of TC-NHEJ complex for DSB repair.
• HTT expression improves DSB repair, genome integrity and phenotypes in HD flies.

## INTRODUCTION

Neurons have high metabolic activity with a robust concomitant generation of reactive oxygen species (ROS) as byproducts of mitochondrial oxidation. These damaging free radicals can escape cellular antioxidant defense systems and trigger oxidative DNA damage in both the nuclear and mitochondrial genomes^1, 2^. In healthy neurons, oxidative DNA damage is efficiently repaired by multiple DNA repair systems to maintain genomic integrity and normal function of neurons^2, 3, 4, 5^, and to prevent the catastrophic premature loss of postmitotic neurons. Yet, defective or impaired DNA repair systems will lead to persistent damage and accumulation of lesions that disrupt normal neuronal gene expression and activate the pro-degenerative DNA damage-response (DDR) signaling. This can culminate in the neuronal dysfunction and cell loss underlying many neurodegenerative diseases^3, 4, 5, 6, 7, 8^.

One of the most lethal forms of DNA damage that occurs in postmitotic neurons is double-stranded breaks (DSBs) which are repaired by nonhomologous end-joining (NHEJ) pathways^3, 5^. The canonical NHEJ pathway involves a core group of NHEJ repair proteins including the Ku70 and Ku80 heterodimer, DNA-dependent protein kinase catalytic subunit (DNA-PKcs), DNA ligase IV (LIG IV), XRCC4, Artemis, PNKP (polynucleotide kinase 3’-phosphtase)^6, 9^, APLF (Aprataxin and PNKP-like factor), and ligase cofactor XLF^3, 5^. DNA-dependent protein kinase (DNA-PK), a complex of the catalytic subunit (DNA-PKcs) and the Ku70-Ku80 heterodimer play principal roles in the most common NHEJ pathway^10, 11^. In response to DSBs, the Ku heterodimer recognizes and binds to the broken DNA ends^11^, followed by recruitment of DNA-PKcs which is activated upon binding to Artemis, a DNA end-processing endonuclease, and finally the DSB repair process is completed by the PNKP-XRCC4-LIG IV complex^12, 13^. Moreover, p53-binding protein (53BP1)^14, 15^ and the Mre11-Rad50-NBS1 (MRN) complex^16, 17^ also play roles in homology dependent DSB repair that is more important in dividing cells but also linked to NHEJ^18^. In response to DSBs, the NHEJ proteins coordinate and repair most of these lesions, thus preserving genome integrity and neuronal function and health.

Recent evidence, including our own studies^19^, suggest that neurons possess added specialized DSB repair mechanism(s) that operate in a dynamic transcriptionally active environment. The transcribing RNA polymerase may recruit DNA repair factors into a multifactorial <u>T</u>ranscription-<u>C</u>oupled DNA <u>R</u>epair (TCR) complex that resolves lesions encountered in the DNA template during transcription^20^. This protection maintains the transcriptionally active genome (euchromatin) and fidelity of the encoded proteins. Although how TCR works to efficiently repair DSBs in neurons remains unclear, the 3,142 reside wildtype huntingtin (wtHTT) acts in a multiprotein complex that repairs DNA *single strand* breaks (SSBs) within transcriptionally active regions in postmitotic neurons^19^. Yet, the wide-spread presence of DNA double strand breaks (DSBs) in Huntington’s disease (HD) brain^21, 22^ implies that a DSB repair mechanism is compromised in HD. In fact, a possible HTT role in DSB repair is suggested due to its interaction with Ku70^21^, an integral component of the DNA-PK complex^11^. Accordingly, overexpressing Ku70 in mouse or *Drosophila* models of HD stimulated NHEJ and decreased neurodegeneration^21, 22^. Furthermore, HTT interacts with NHEJ protein PNKP^19^ and XRCC4-LIG IV complex^6, 23^.

To test the potential role of HTT in NHEJ repair of DSBs during transcription in neurons, we here combined biochemistry with computation and molecular cell biology on HTT interactions. We found that wtHTT and chromatin remodeling protein BRG1 scaffold an efficient transcription-coupled non-homologous end-joining (TC-NHEJ) complex in brain. The results suggest that this multifactorial complex repairs DSBs that routinely occur in transcribed gene-rich areas in neurons. We furthermore find that mutant HTT (mHTT) in the TC-NHEJ complex compromises its DSB repair activity to undermine the functional integrity of the neuronal genome consistent with onset of HD phenotypes. Moreover, our functional studies revealed that mHTT expression in *Drosophila* impairs DSB repair; however, increasing HTT and TC-NHEJ activity can mitigate neurotoxicity including the motor defects and reduced longevity in mutant HD flies.

## RESULTS

### HTT and BRG1 assemble into a transcription-coupled NHEJ complex

To evaluate whether HTT is a component of an active NHEJ complex, we examined whether wtHTT interacts with NHEJ components. We first immunoprecipitated HTT from wild-type neuroblastoma SH-SY5Y cell-derived nuclear extracts (NEs) with an anti-HTT antibody and analyzed the HTT immunocomplexes (ICs) by mass spectrometry (MS). MS analyses of the HTT ICs revealed a consistent presence of key NHEJ factors including Ku70 (XRCC6), Ku80 (XRCC5), XRCC4, DNA-PKcs (PRKDC), PNKP, DNA Ligase IV (LIG IV), 53BP1 (p53-binding protein 1), MRE11 and RAD50, and several RNA-binding/processing factors (Figure 1A). MS analysis furthermore found chromatin remodeling protein Brahma-Related Gene 1 (BRG1) and other related chromatin remodeling proteins in HTT ICs (Figures 1A to 1C). BRG1, also known as SMARCA4 (SWI/SNF-related, Matrix-associated, Actin-dependent Regulator of Chromatin, subfamily A, member 4) is the central ATPase subunit of the SWI/SNF-like chromatin remodeling complexes with critical roles in regulating chromatin dynamics^24, 25^ as well as DNA repair^26, 27, 28^. Finding BRG1 and several key NHEJ proteins provided an additional indication that HTT might be integral to the assembly of a functional multifactorial TC-NHEJ complex in the nervous system (Figure 1C). HTT’s association with BRG1 furthermore suggests that HTT might be intimately related to the chromatin assembly/remodeling processes in the nervous system.

**Figure. 1.**
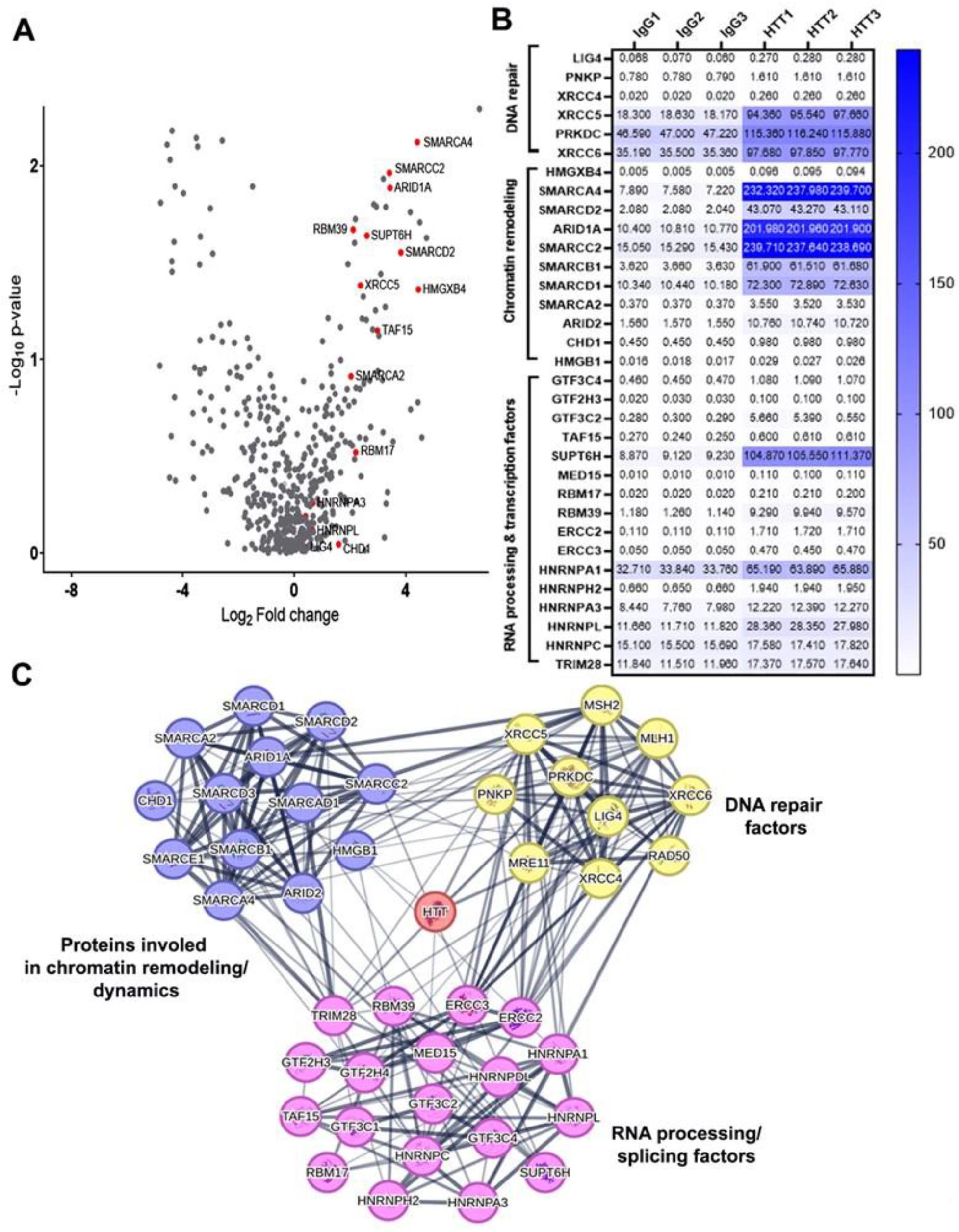
Nuclear HTT interacts with DNA repair, chromatin remodeling, RNA processing, and transcription factor proteins. **(A)** HTT interacting proteins. Volcano plot displays HTT interacting proteins (right) and non-specific control IgG proteins (left). Each black dot represents a protein. and the Red points indicate significant HTT interaction proteins in DNA repair, chromatin remodeling and dynamics, RNA processing and splicing, and transcription. The nuclear fraction from SH-SY5Y cells was co-immunoprecipitated with either HTT or control IgG antibodies, followed by mass spectrometry (MS) analysis. **(B)** Heat map showing the relative interaction of either HTT or control IgG with various proteins in the nuclear fractions. Respective fold change values (x10^5^, right) are included with white (low) to blue (high) color map for increasingly stronger interaction from three biological replicates (n=3). **(C)** STRING database-based network of HTT-interacting proteins reveals HTT interactions with DNA repair, chromatin remodeling, RNA processing, and transcription proteins. In this network, each node represents a protein, while each edge indicates a physical and/or functional interaction based on STRING data.

To further characterize the putative TC-NHEJ complex and to validate the MS findings, we immunoprecipitated HTT from the NEs isolated from 3-month-old wildtype C57BL/6 mouse brain tissue (Cortex) and analyzed the resultant ICs for the presence of key NHEJ factors by western blotting. The results not only revealed Ku70 as described earlier^21^, but also confirmed the presence of BRG1 along with several established NHEJ factors including DNA-PKcs, PNKP, XRCC4, DNA LIG IV, and RNA polymerase II large subunit A (POLR2A) in HTT ICs (Figure 2A). The presence of POLR2A, Cockayne Syndrome protein B (CSB), and transcription elongation factor TFIIS, which are key proteins necessary for initiating and regulating nuclear TCR^29, 30, 31^, in the HTT ICs (Figure 2A) supports HTT as a fundamental constituent of a transcription-coupled NHEJ (TC-NHEJ) complex, with BRG1 as a novel component of this complex.

**Figure 2.**
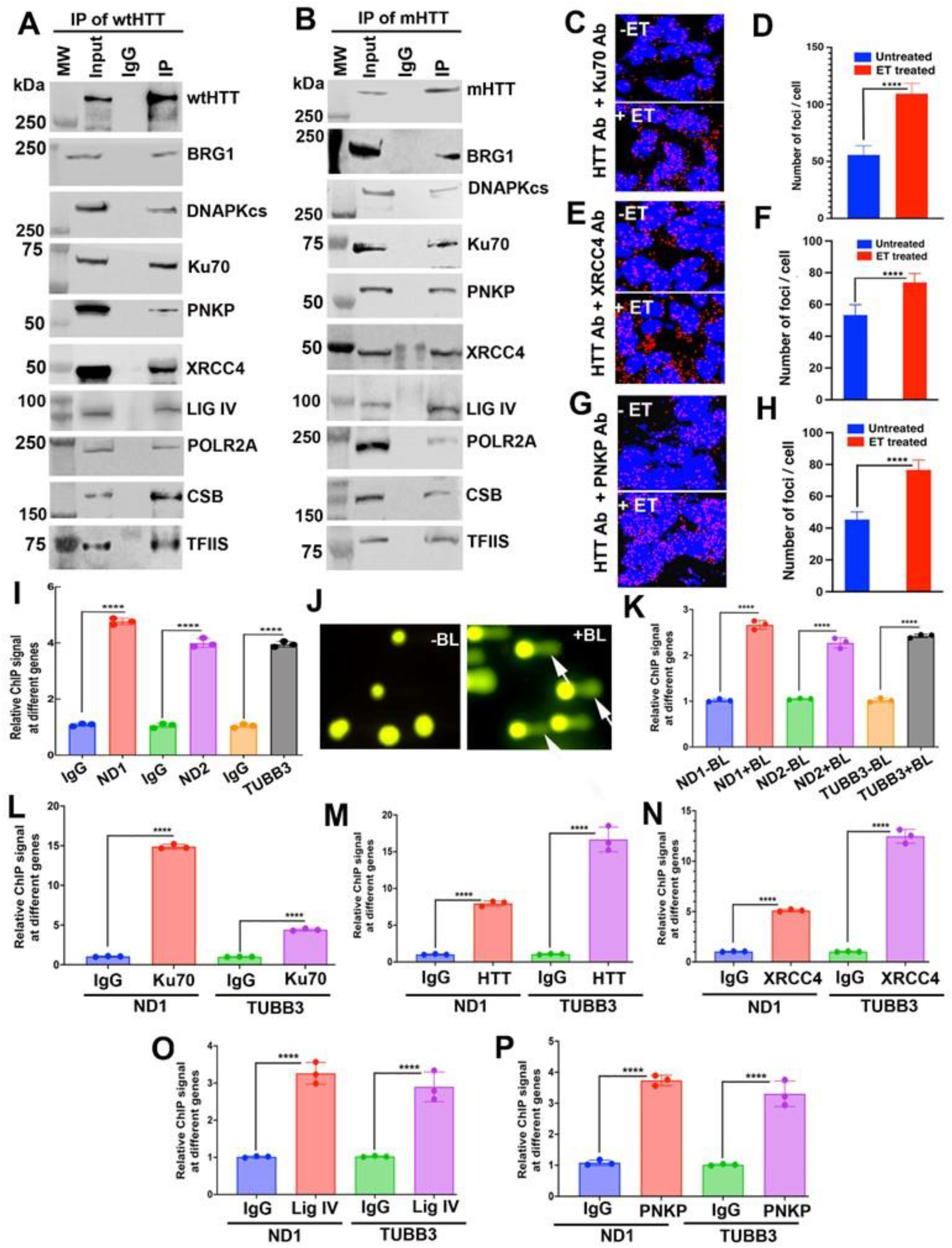
An implicated huntingtin (HTT)-assembled transcription-coupled non- homologous end joining (TC-NHEJ) repair complex. **(A)** HTT immunocomplexes (ICs) analyzed by western blots (WBs) from nuclear protein extracts (NEs) isolated from 3-month-old wild-type C57BL/6 mouse brain. HTT was immunoprecipitated (IP’d) from NEs with an anti-HTT monoclonal antibody (Ab) (MAB2710, Millipore-Sigma). Immunocomplexes (ICs) tested for BRG1 and TC- NHEJ factors DNA-PKcs, Ku70, PNKP, XRCC4, LIG IV, POLR2A, CSB, and TFIIS in HTT ICs. Lane 1; protein molecular weight marker; lane 2: Input; lane 3: IgG control IP; lane 4: IP of HTT with anti-HTT Ab. **(B)** mHTT WBs IP’d from NEs isolated from 3-month-old HD homozygous zQ175 transgenic mouse brain with an anti-HTT Ab (MAB2710, Millipore-Sigma). mHTT ICs were analyzed by WBs for BRG1, and TC-NHEJ factors. Lane 1; protein molecular weight marker; lane 2: Input; lane 3: IgG control IP; lane 4: IP of HTT with an anti-HTT Ab. **(C-G)** Proximity ligation assay (PLA) for HTT and NHEJ factors in wild-type SH-SY5Y and in SH-SY5Y cells after treating the cells with DNA damaging agent etoposide (ET; 15µM, for 30 minutes). Generation of red fluorescence indicates representative positive protein-protein interactions. Nuclei stained with DAPI. (C) HTT interaction with Ku70 changes in response to increased DSBs. PLA was performed on SH-SY5Y cells with anti-HTT rabbit monoclonal Ab (5656; Cell Signaling) and anti-Ku70 mouse monoclonal Ab (SC-5309; Santa Cruz) before and after treating cells with ET (-ET and + ET respectively). **(D)** HTT and Ku70 relative interactions by PLA signals in control untreated (-ET) and ET- treated (+ ET) cells. Data represent mean ± SD, ****p<0.0001. **(E)** HTT interactions with XRCC4 in response to increased DSBs. PLA was performed with rabbit monoclonal anti-HTT Ab (5656; Cell Signaling) and anti-XRCC4 mouse monoclonal Ab (SC-271087; Santa Cruz) before and after treating the SH-SY5Y cells with ET (-ET and + ET respectively). **(F)** HTT and XRCC4 relative interactions assessed by relative PLA signals in control (- ET) cells and ET-treated (+ ET) cells. Data represent mean ± SD, ****p<0.0001. **(G)** HTT and PNKP interactions in response to increased DNA damage. PLA performed with anti-HTT mouse monoclonal Ab (MAB2170; Millipore-Sigma) and anti-PNKP rabbit polyclonal Ab (MBP-1-A7257; Novus) before and after treating SH-SY5Y cells with ET (-ET and + ET respectively). **(H)** HTT and PNKP interactions assessed by relative PLA signals in control (- ET) cells and ET-treated (+ ET) cells. Data represent mean ± SD, ****p<0.0001. **(I)** HTT association with genome. Chromatin immunoprecipitation (ChIP) performed to assess association/interaction of HTT with genome in untreated SH-SY5Y cells in a stress-free condition. After ChIP, DNA fragments purified from the cross-linked protein-DNA entities and the purified genomic DNA fragments analyzed by real-time quantitative PCR (qPCR). Relative ChIP values measured with respect to control IgG. Data represent mean ± SEM, ****p<0.0001. **(J)** DSBs measured by neutral comet analysis. SH-SY5Y cells were treated with DNA damaging agent bleomycin (BL; 5µg/mL, for 30 minutes), and subjected to neutral comet analysis to detect DSBs before (-BL) and after BL treatments (+ BL). Increased comet tail moments after BL treatments indicate presence of DSBs (shown with arrows; right panel). **(K)** HTT association with genome before and after inducing DSBs. SH-SY5Y cells treated with BL for 30 minutes to induce DSBs, and ChIP performed on control untreated cells and BL-treated cells. Relative ChIP values measured after normalization to control IgG. Data represent mean ± SEM, ****p<0.0001. (**L-P**) Sequential ChIP (ChIP-re-ChIP) analysis was performed to determine whether HTT and NHEJ factors co-occupy the same genomic DNA loci *in vivo*. **(L)** HTT-cross-linked genomic DNA. DNA fragments were isolated from the wildtype C57BL/6 mouse brain, and the DNA-protein complexes IP’d with an anti-HTT Ab (MAB2170; Millipore-Sigma), followed by a second immunoprecipitation (IP) of the HTT ICs with either an anti-Ku70 or an anti-IgG Abs. Genomic DNA fragments were isolated from the final Ku70 ICs and IgG ICs, and genomic DNA segments (∼250 bp) encompassing Neurod1 (ND1) or Tubulin Beta 3 Class III (Tubb3) genomic regions were amplified by quantitative PCR (qPCR) using specific primers and PCR products were quantified by qPCR. Relative ChIP values were measured after normalization to control IgG. Data represent mean ± SEM, ****p<0.0001. **(M)** Ku70-bound genomic DNA. DNA fragments were IP’d with an anti-Ku70 Ab (SC- 5309; Santa Cruz), followed by a second IP of the Ku70 ICs with an anti-HTT (MAB2170; Millipore-Sigma) or an anti-IgG Abs. DNA isolated from the final HTT ICs and IgG ICs, and genomic DNA segments (∼250 bp) encompassing Neurod1 or Tubb3 genes was amplified using specific primers and the PCR products quantified. Relative ChIP values were measured after normalization to control IgG. Data represent mean ± SEM, ****p<0.0001. **(N)** HTT-DNA complexes from anti-HTT and anti-XRCC4 Abs. Putative HTT-DNA complexes were IP’d with an anti-HTT Ab (MAB2170; Millipore-Sigma), followed by a second IP of the HTT ICs with either an anti-XRCC4 (SC-271087; Santa Cruz) or an anti-IgG Abs. The DNA fragments were isolated from XRCC4 ICs and IgG ICs, and genomic DNA segments (∼250 bp) encompassing Neurod1 or Tubb3 genes amplified, and PCR products analyzed by qPCR. Relative ChIP values measured after normalization to IgG. Data represent mean ± SEM, ****p<0.0001. **(O)** HTT-DNA complexes from anti-HTT and anti-DNA ligase IV Abs. Putative HTT- DNA complexes were IP’d with an anti-HTT Ab (MAB2170; Millipore-Sigma), followed by a second IP of the HTT ICs with either an anti-DNA ligase IV (14649; Cell Signaling) or an anti-IgG Abs. DNA isolated from the ligase IV ICs and IgG ICs, and genomic DNA segments encompassing Neurod1 or Tubb3 genes were amplified by qPCR and the PCR products analyzed. Relative ChIP values measured after normalization to control IgG. Data represent mean ± SEM, ****p<0.0001. **(P)** HTT-DNA complexes from anti-HTT and anti-PNKP Abs Putative HTT-bound DNA fragments were IP’d with an anti-HTT Ab (MAB2170; Millipore-Sigma), followed by a second IP of the HTT ICs with either an anti-PNKP Ab (MBP-1-A7257; Novus) or an anti-IgG Abs. DNA isolated from PNKP ICs and IgG ICs, and genomic segments (∼250 bp) encompassing Neurod1 or Tubb3 genes were amplified using specific primers and the PCR products quantified. Relative ChIP values measured after normalization to control IgG. Data represent mean ± SEM, ****p<0.0001.

We next tested whether the expanded polyQ sequences in mutant HTT (mHTT) interferes with the integrity of this complex. We isolated NEs from 3-month-old Q175 homozygous HD transgenic mouse cortex (CTX)^32^, immunoprecipitated mHTT with the anti-HTT Ab and subjected the isolated complexes to western blotting. The data revealed the same NHEJ factors in the HTT ICs (Figure 2B), suggesting that mHTT is present in the TC-NHEJ complex and does not compromise its composition *in vivo*. Because Ku70 was detected in the HTT ICs, we next performed a reciprocal immunoprecipitation experiment using an anti-Ku70 Ab on the same NEs, and western blot analyses confirmed that HTT, BRG1, POLR2A, and several other NHEJ factors co-precipitated with Ku70 (Supplementary figure 1A). Likewise, immunoprecipitation of DNA-PKcs with an anti-DNA-PKcs Ab or XRCC4 with an anti-XRCC4 Ab yielded qualitatively consistent results, detecting HTT, BRG1, POLR2A and several NHEJ proteins (Supplementary Figure 1B and 1C, respectively). To examine the specificity of these interactions, we analyzed the ICs for the presence of apurinic-apyrimidinic endonuclease 1 (APE1), an abundant critical DNA base excision repair (BER) enzyme that works independently of PNKP-mediated repair^19^. APE1 was not detected in the ICs (Supplementary figures 1A to 1C), supporting the specificity and selectivity of the observed protein interactions *in vivo*.

To further examine HTT interaction with key NHEJ proteins, we performed a proximity ligation assay (PLAs) in SH-SY5Y cells. Consistent with our immunological studies, PLA results indicate close spatial interactions between HTT and NHEJ factors including Ku70, XRCC4 and PNKP. Notably, these interactions were markedly increased following induction of DSBs by treating the cells with etoposide (ET; 15µM; 30 minutes) (Figure 2C to 2H). PLAs also detected notable interactions between HTT and Ku80, DNA-PKcs, CSB, and LIG IV, which increased when DSBs were introduced (Supplementary figures 1D to 1K). Since PNKP and HTT are present in mitochondria and can regulate DNA repair, fluorescent signaling outside nuclei are presumably from mitochondrial complexes. Importantly, HTT interaction with PARP1 was unaltered when DNA damage was induced with ET (Supplementary figure 1L and 1M), suggesting that PARP1 may be constitutively associated with the genome, in contrast to Ku70, Ku80, DNA-PKcs, XRCC4, LIG IV and PNKP which are recruited more into the TC-NHEJ complex in response to increased DNA lesions.

We reasoned that if HTT is a functional DSB DNA repair complex component, it would physically associate with the genome in a manner that would increase at sites of new DSBs. We therefore performed chromatin immunoprecipitation (ChIP) assays with anti-HTT Ab to examine the HTT genomic interaction as a function of DNA damage in SH-SY5Y cells. The ChIP analysis showed a pronounced HTT-DNA interaction in SH-SY5Y cells at several sites in the genome (Figure 2I). To test whether HTT recruitment is enhanced in response to increased DNA damage, we introduced DSBs in genomic DNA by treating the cells with bleomycin (BL; 5µg/ml), which induces DSBs^33^. Neutral comet analysis^34^ revealed BL treatment increased DSBs in SH-SY5Y cells (Figure 2J; arrows). ChIP analysis showed 2- to 3-fold increased HTT interaction with the genome upon induction of DSBs (Figure 2K). As a positive control, we assessed DNA binding by PNKP, an established TC-NHEJ repair enzyme^6, 9^ before and after inducing DSBs. We found an increased association of PNKP with the genome when DSBs were induced (Supplementary figure 1N).

Given that HTT interacted with NHEJ components, and they temporally co-localize to the genome, we investigated whether HTT could be a core TC-NHEJ complex component *in vivo*. We therefore used sequential ChIP (ChIP-re-ChIP) analysis^19, 35^ to further test the co-occupancy of HTT and interacting NHEJ proteins with the DNA. We sequentially immunoprecipitated protein-DNA complexes in wildtype C57BL/6 mouse (3-months-old) cortical tissues, initially with the anti-HTT Ab followed by second immunoprecipitation using an anti-Ku70 Ab representative of the NHEJ complex. Quantitative PCR (qPCR) analysis of select genomic sites established that HTT and Ku70 co-localized, consistent with recruitment of the NHEJ complexes harboring HTT (Figure 2L). Reciprocal ChIP-re-ChIP studies sequentially targeting Ku70 and HTT respectively, validated formation of the complexes (Figure 2M). Subsequent ChIP-re-ChIP experiments centered on additional NHEJ components and confirmed that XRCC4, LIG IV, and PNKP also co-localized with HTT at specific genomic sites (Figure 2N-2P). These combined observations indicate that HTT associates with several NHEJ components on DNA, consistent with a functional role in transcription-coupled DSB repair in the central nervous system.

### BRG1 directly interacts with HTT

Chromatin remodeler BRG1 modifies DNA-histone contacts to alter chromatin architecture and modulation of transcription^24, 25^, and BRG1 also acts in DNA repair and genome integrity^26, 27, 28^. Our observation that BRG1 is present in the HTT ICs prompted us to test whether BRG1 is a part of the HTT-associated TC-NHEJ complex, and if present to characterize the nature of the HTT interaction. Wildtype C57BL/6 mouse brain NEs were immunoprecipitated with an anti-BRG1 Ab (NB100-2594; Novus) under stringent conditions to control for non-specific protein interactions. Western blotting of BRG1 and the co-precipitated proteins detected HTT along with several NHEJ factors (Figure 3A), corroborated earlier MS findings. These data were further validated by using epitope-tagged proteins in co-immunoprecipitation studies. FLAG-wtHTT-Q19 (FLAG-tagged full-length wtHTT encoding 19 glutamine) and Myc-BRG1 (Myc-tagged BRG1) were co-expressed in SH-SY5Y cells, followed by isolation of NEs and immunoprecipitation of FLAG-tagged HTT with an anti-FLAG Ab, and the presence of Myc-BRG1 assayed. The results show BRG1 in the FLAG ICs (Figures 3B and 3C). We then performed PLA on SH-SY5Y cells to assess HTT interaction with BRG1. PLA showed interactions between HTT and BRG1 under stress-free conditions, which greatly increased upon induction of DSBs by treatment with 15µM etoposide for 30 minutes (Figures 3D and 3E), in keeping with BRG1-HTT acting in DSB repair^26, 27, 28^.

**Figure 3.**
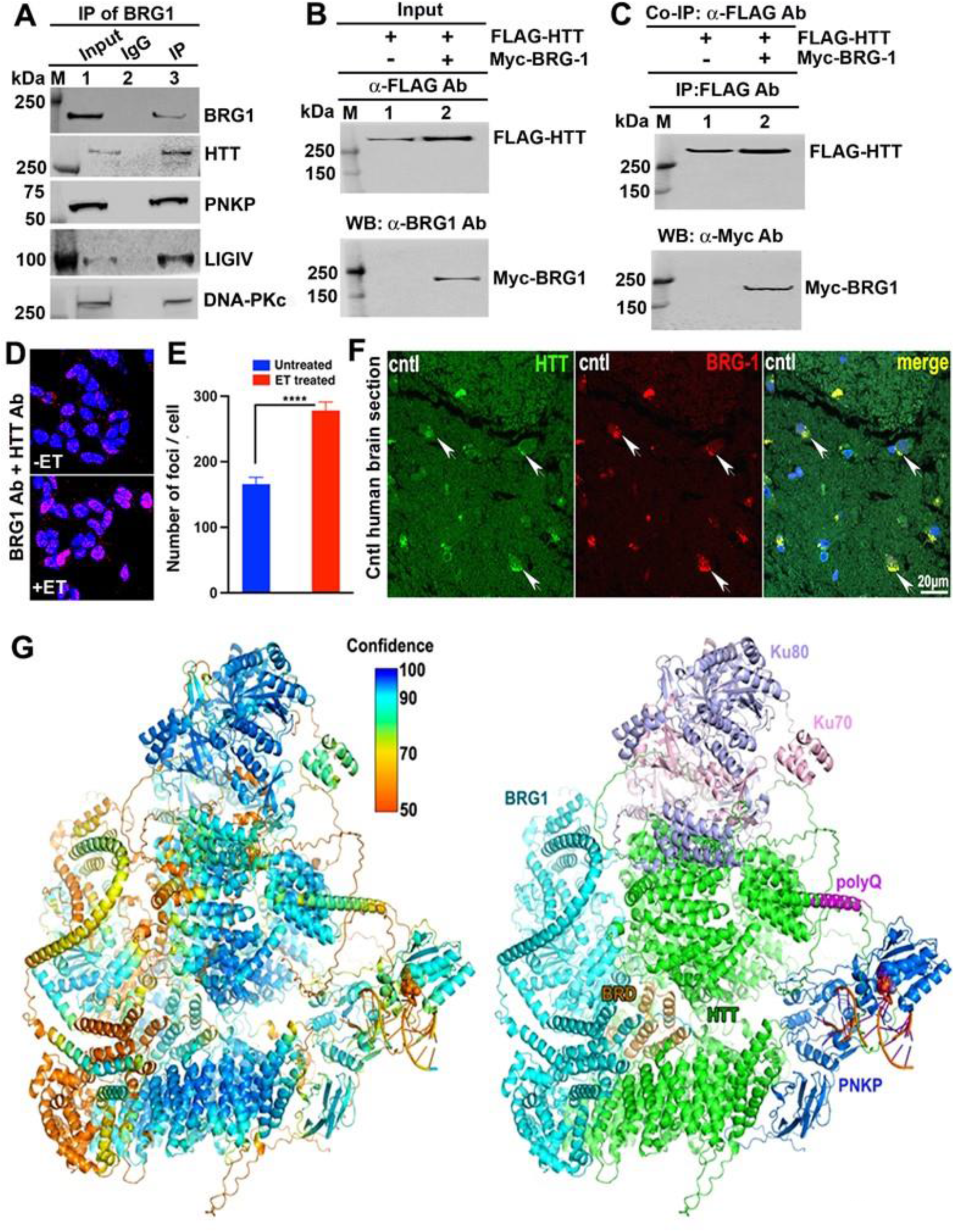
BRG1 chromatin remodeler associates with HTT *in vivo* as a key component of an HTT-assembled TC-NHEJ complex. **(A)** BRG1 ICs contain HTT and NHEJ components. NEs isolated from 3-month-old wild-type C57BL/6 mouse brain tissue (cortex) and BRG1 were IP’d from the NEs with an anti-BRG1 Ab (NB100-2594; Novus). Resulting BRG1 ICs were analyzed by WBs to detect BRG1, HTT and associated NHEJ components in BRG1 ICs. Lane M; protein molecular weight marker; lane 1: Input; lane 2: IgG control; lane 3: IP of BRG1 with anti-BRG1 Ab. **(B)** FLAG-HTT and Myc-BRG1 association. Plasmids p-FLAG-HTT-Q24 and p-Myc- BRG1, expressing FLAG-tagged wtHTT-Q24 and Myc-tagged human BRG1, respectively, were co-transfected into SH-SY5Y cells, cells harvested and NEs isolated 48 hours post-transfection. Expressions of FLAG-HTT and Myc-BRG1 were analyzed by WBs with anti-FLAG Ab (F3165; Millipore-Sigma) and anti-Myc 9E10 Ab (SC- 40; Santa Cruz) respectively. **(C)** Myc-tagged BRG1 and FLAG-tagged human HTT-Q24 association. Plasmids p- FLAG-HTT-Q24 and p-Myc-BRG1, expressing FLAG-tagged human HTT-Q24 and Myc-tagged human BRG1, respectively, were co-transfected into SH-SY5Y cells, NEs isolated from the cells 48 hours post-transfection. FLAG-tagged HTT-Q24 were IP’d from the NEs with an anti-FLAG Ab (F3165; Millipore-Sigma), and FLAG ICs analyzed by WBs with an anti-Myc 9E10 Ab (SC-40; Santa Cruz) to detect Myc-tagged BRG1 in FLAG ICs. **(D)** HTT and BRG1 proximity before and after ET induced DNA damage. Proximity ligation assays (PLA) were performed to assess possible interaction of HTT and BRG1 before (upper panel) and after (lower panel) inducing DNA damage by treating cells with ET (300 ng/mL, for 30 minutes). Reconstitution of red fluorescence indicates interaction of HTT with BRG1. Nuclei stained with DAPI. **(E)** Proximity of endogenous HTT with BRG1. Relative signals show PLA interactions of endogenous HTT with BRG1 in control untreated SH-SY5Y cells (Cntl; -ET), and in SH-SY5Y cells treated with ET (+ET; 300 ng/mL for 30 minutes). Data represent means ± SD, ****p<0.0001. **(F)** HTT and BRG1 co-localize in human brain. Postmortem human brain sections (striatum) analyzed by co-immunostaining with an anti-HTT mouse monoclonal Ab (MAB2170; Millipore-Sigma) and anti-BRG1 rabbit polyclonal Ab (3508; Cell Signaling) and sections analyzed by confocal microscopy. Nuclei stained with DAPI, and co-localization of HTT (green fluorescence) with BRG1 (red fluorescence) appear as yellow fluorescence (shown by arrows). **(G)** Predicted HTT-mediated protein assembly with BRG1, PNKP-DNA, and Ku70/80. AlphaFold3 (AF3) predicted complex is shown with color-coded confidence levels (left) and labeled protein chains (right). AF3 places HTT (green) in the core of an NHEJ protein complex (right). Ku70/80 (purple/pink, top) has a significant interface with HTT, which is sandwiched between BRG1 (cyan) and PNKP (blue) that neighbors the PolyQ extension (purple). The BRG1 Bromodomain (BRD: brown) fits into the HTT central cleft. HTT interfaces with BRG1 BRD and Ku70/80 have the highest AF3 confidence levels (left).

BRG1 interacts with proteins involved in DNA repair and DNA-damage-response (DDR) including PARP1, BRCA1 and p53^36^, which suggest BRG1 participates in DSB repair. Since BRG1 interacts with HTT, we sought to identify NHEJ proteins that may directly interact/associate with BRG1 and HTT. First, to identify the NHEJ factor(s) that BRG1 interacts with in the TC-NHEJ complex, we expressed FLAG-tagged BRG1 (FLAG-BRG1) in SH-SY5Y cells, isolated the NEs and immunoprecipitated FLAG-BRG1 from the NEs with an anti-FLAG Ab. Assessment of the co-immunoprecipitated proteins by western blotting showed the presence of DNA-PKcs, and HTT in the FLAG ICs but not Ku70, Ku80, XRCC4, LIG IV or PNKP (Supplementary figure 2A) in the FLAG ICs, suggesting that BRG1 selectively and specifically interacts with HTT and DNA-PKcs and these associations might help assemble the functional macromolecular TC-NHEJ complex. Since BRG1 interacts with HTT we further tested whether HTT also interacts with DNA-PK and other essential NHEJ factors. To examine this, we expressed FLAG-HTT-Q19 in SH- SY5Y cells, isolated NEs and immunoprecipitated HTT from the NEs with the anti-FLAG Ab under stringent IP condition designed to limit immunoprecipitation of proteins that are not interacting/associating with HTT. Subsequent western blot analysis of the FLAG ICs revealed the presence of key NHEJ factors included BRG1, POLR2A, DNA-PKcs, Ku70, Ku80, XRCC4, PNKP, and LIG IV (Supplementary figure 2B), suggesting that endogenous HTT interacts/associates with BRG1 and several key NHEJ proteins. To further assess the possible association of BRG1 with NHEJ proteins we performed PLAs in SH-SY5Y cells. The PLA findings corroborate the immunoprecipitation results by showing that BRG1 interacts with the NHEJ proteins, (Supplementary figure 2C). Collectively, these data suggest that both HTT and BRG1 are part of the same TC-NHEJ complex. To further test the implications of these combined cellular results *in vivo*, confocal image analysis of postmortem brain sections (striatum; 56-year-old) from control, non-diseased individuals revealed a pronounced co-localization of HTT and BRG1 (Figure 2F; arrows), indicating potential *in vivo* interactions.

Although the large size of HTT and BRG1 make computational modeling challenging, we reasoned that a computational structure prediction by AlphaFold 3 (AF3) could provide an independent assessment of the potential for their structural interaction^37^. The objective AF3 computational structure supported HTT-mediated protein assembly with BRG1, PNKP-DNA, and Ku70/80 (Figure 2G). Notably, AF3 places HTT in the core of this complex with HTT being sandwiched between BRG1 and PNKP (Figure 2G). BRG1 Bromodomain (BRD) is tucked into the HTT central cleft with high confidence levels consistent with the cell biology. Collectively, these cell culture, computational, and tissue data suggest that HTT and BRG1 are integral components of a functional neuronal TC-NHEJ complex.

### HTT facilitates DSB repair to maintain neuronal genome integrity

Although many papers suggest that increasing wtHTT levels improves neuronal resistance to degeneration^38, 39, 40, 41, 42, 43, 44, 45, 46, 47^, the mechanism by which the wtHTT protein may provide neuroprotection against genotoxic agents remains unknown. We reasoned that HTT-mediated assembly of the TC-NHEJ complex for repair of DSBs, which are the most lethal DNA lesions in postmitotic cells like neurons, could be a significant mechanism for neuroprotection in the central nervous system.

To test whether wtHTT facilitates DSB repair, we modulated HTT levels in SH-SY5Y cells by over-expressing exogenous wtHTT [HTT-OE], or by suppressing wtHTT levels using RNA interference [HTT-KD], respectively. Neither overexpression nor knock-down of HTT in SH- SY5Y cells interfered with expression levels of the key NHEJ proteins including Ku70, Ku80, PNKP, and XRCC4 (Figure 4A). In response to DSBs, several DNA repair proteins including histone H2AX and 53BP1 (p53-binding protein 1) are phosphorylated and rapidly recruited to damage sites, which is detected as discrete puncta by immunostaining using γH2AX- or 53BP1- specific Abs^48^. Immunostaining with an anti-53BP1 Ab under stress-free conditions showed widespread presence of 53BP1-positive nuclear foci in the HTT-KD cells (Figure 4B; upper panel; arrows). These were absent in control cells (Figure 4B; lower panel), suggesting that accumulated DSBs remain largely unrepaired in HTT-deficient cells.

**Figure 4.**
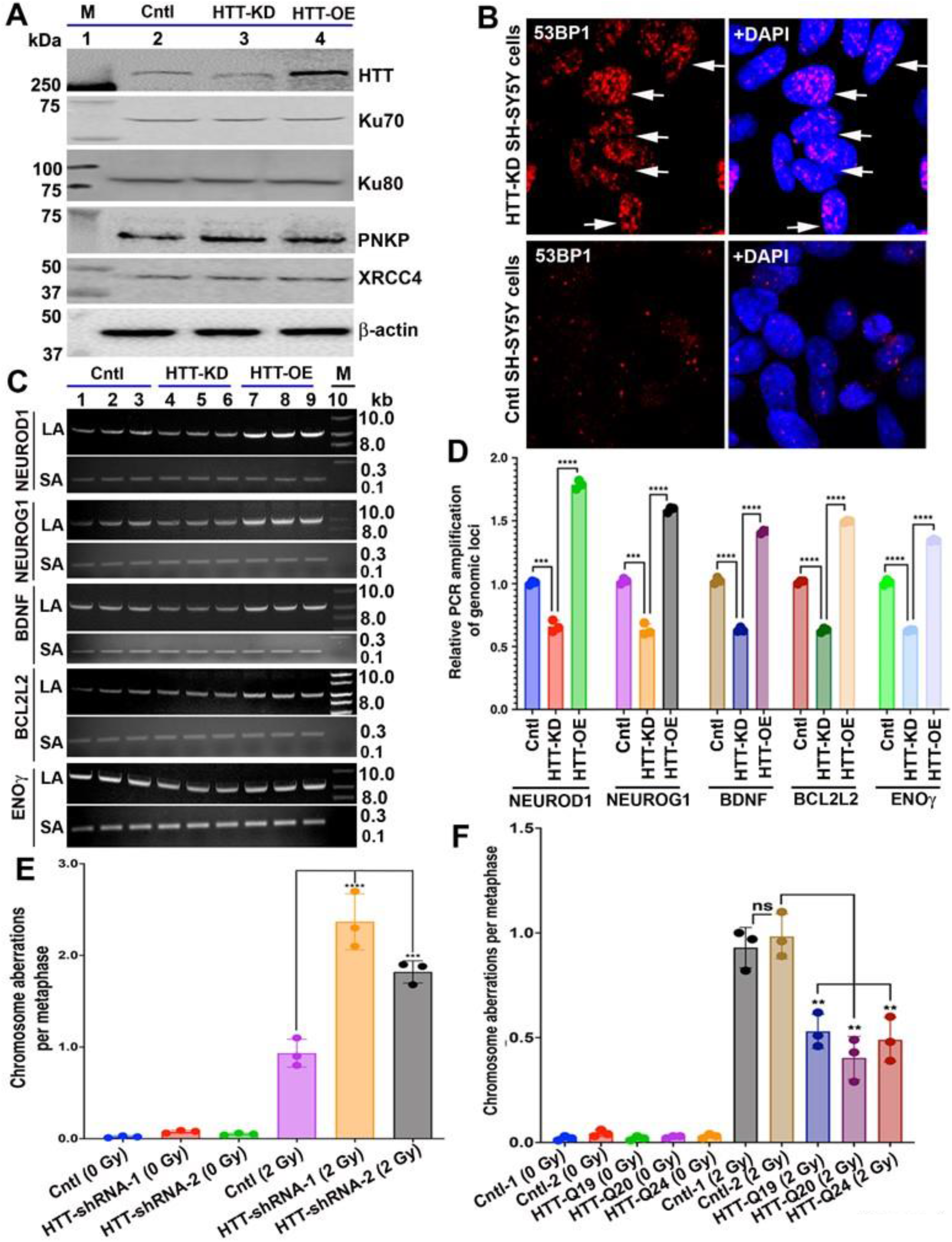
Wildtype HTT stimulates DSB repair to maintain genome integrity. **(A)** Experimental HTT association with Ku70, Ku80, PNKP, and XRCC4. Total proteins were extracted from control SH-SY5Y cells (Cntl; lane 2), SH-SY5Y cells either expressing HTT-RNAi (HTT-KD cells; lane 3) or full-length human wtHTT cDNA (HTT-OE cells; lane 4). Protein extracts were analyzed by WBs to detect HTT, Ku70, Ku80, PNKP, and XRCC4 levels; β-actin was used as loading control. Lane 1: Protein molecular weight marker in kDa. **(B)** HTT-depletion increases double strand break (DSB) foci. HTT-knocked down (HTT- KD) SH-SY5Y cells (upper panels) and control wildtype untreated SH-SY5Y cells (lower panels) were analyzed by immunostaining with an anti-53BP1 Ab (4937; Cell Signaling) to measure DSBs in the nuclei. The 53BP1-positive DSBs in genomic DNA (red puncta) within the nuclei (blue) of HTT-KD cells (upper panel) shown by arrows. Similar discrete nuclear puncta not detected in control cell nuclei (Lower panel). Nuclei stained with DAPI (blue). **(C)** HTT-depletion increases measured genomic DNA damage. Genomic DNA isolated from control SH-SY5Y cells (Cntl; lanes 1 to 3), SH-SY5Y cells expressing HTT- RNAi (HTT-KD cells; lanes 4 to 6) or SH-SY5Y cells overexpressing full-length wtHTT (HTT-OE cells; lanes 7 to 9), and presence of DNA damage/lesions in the genome were assessed by LA-QPCR analyses. Genomic DNA segments (8 to 10 kb segments) from various genome regions encompassing the NEUROD1, NEUROG1, BDNF, BCL2L2, or ENOLASE gamma (ENOγ) genes PCR were amplified using specific primers, and the PCR products analyzed on agarose gel, and intensity of the DNA bands quantified. LA denotes long amplicon (8.0 to 9.0 kb); SA denotes short amplicon (0.2 to 0.3 kb); Lane 10: 1-kb DNA ladder. **(D)** HTT-depletion significantly increases DNA damage in multiple genomic regions. Relative levels of DNA damage in various genomic DNA regions encompassing the NEUROD1, NEUROG1, BDNF, BCL2L2, or ENOγ genes quantified in control cells (Cntl), and cells expressing HTT-RNAi (HTT-KD) or full-length HTT-cDNA (HTT- OE). Data represents Mean ± SD. ***p<0.001; ****p<0.0001. **(E)** HTT-depletion increases chromosome aberrations from ionizing radiation. Control SH- SY5Y cells and HTT-deficient SH-SY5Y cells (expressing two independent shRNA; HTT-shRNA-1 and HTT-shRNA-2) were exposed to ionization radiation (2 Gray units) and chromosome aberration at metaphase scored before and after irradiation from three independent experiments. For each experiment chromosome aberrations from 100 metaphases scored. (Mean ± SD. ***p<0.001; ****p<0.0001). **(F)** HTT PolyQ length and chromosome aberrations from ionizing radiation. Control wildtype SH-SY5Y cells (SH-SY5Y-1 and SH-SY5Y-2 cells) and SH-SY5Y cells overexpressing exogenous FLAG-tagged wtHTT encoding 19, 20 or 24 glutamines (SH-SY5Y-HTT-Q19, SH-SY5Y-HTT-Q20, and SH-SY5Y-Q24) were exposed to ionization radiation (2 Gray units) and chromosome aberration at metaphase scored before and after irradiation from three independent experiments. For each experiment chromosome aberrations from 100 metaphases scored. (Mean ± SD. **p<0.01); ns denotes not significant.

To further test the notion that HTT deficiency hampers DNA repair, we used Long Amplicon Quantitative PCR (LA-QPCR) analysis^19^ to quantify the amount of DNA damage in the HTT-KD cells compared with the control cells. LA-QPCR analyses of genomic DNA from HTT- KD cells revealed that PCR amplification of various genome segments (NEUROD1, NEUROG1, BDNF, BCL2L2, and ENOLASE2 gamma [ENO2γ]) were significantly diminished compared with the amplification efficiencies of the same genome segments from control cells (Figures 4C; lanes 4-6 vs. lanes 1-3, and figure 4D). These data reinforce our observation that HTT-KD cells exhibit a compromised capacity to resolve DSBs, culminating in persistent DNA damage. We next tested whether increasing wtHTT expression in neuronal cells would enhance DNA repair and improve overall genome integrity. wtHTT was overexpressed in SH-SY5Y cells, genomic DNA isolated and DNA damage quantified in HTT-OE cells and control cells by LA-QPCR analysis. PCR amplification yielded significantly more amplicons for the genomic regions examined with template DNA isolated from the HTT-OE cells as compared with control cells (Figures 4C; lanes 7-9 vs. lanes 1-3, and Figure 4D). This measurement indicated reduced DNA damage in the HTT- OE cells relative to controls as evidenced by more efficient and robust DSB repair.

To more directly examine how HTT promotes DSB repair and genome integrity, we induced DSBs in the genome by irradiating the HTT-KD and control SH-SY5Y cells with ionizing radiation (IR; 2 Gray units) and quantification of relative chromosome aberrations. Control cells showed significant chromosome aberrations when exposed to IR, whilst HTT-KD cells exhibited even higher (2-fold increase) chromosome aberrations compared with control cells exposed to the same dose of IR (Figure 4E). Together, these data suggest that HTT-deficient cells are susceptible to more DNA lesion accumulation and increased chromosome aberrations and genome instability when exposed to damaging agents. Conversely, SH-SY5Y cells overexpressing exogenous wtHTT exhibited significant resistance to chromosome aberrations compared with control cells when exposed to the same dose of IR under identical experimental conditions (Figure 4F).

Consistent with these results, neutral comet analyses reveal more DSBs in the HTT-KD cells compared with control cells under stress-free conditions (Supplementary figure 3A and 3B; arrows). To test whether HTT-deficiency impairs NHEJ-mediated DSB repair, we next measured NHEJ-mediated DSB repair efficiency in HTT-KD cells and control cells by using a GFP method described previously^49^. The NHEJ-mediated error-free repair of I-SceI sites after DSB induction results in expression of functional GFP, and therefore the percentage of GFP-positive cells provides a relative measure of NHEJ-mediated DSB repair. Cells positive for GFP were observed at chromosome 1 site A (Chr1-A) (transcribing region), as compared to Chr1-B (non-transcribing region), with higher drop in NHEJ-mediated repair (>70%) in Chr1-A as compared to Chr1-B in HTT-KD cells, suggesting efficiency of NHEJ-mediated DSB repair is markedly compromised in HTT-KD cells compared with control cells (Supplemental figure 3C). These findings strongly suggest that effective NHEJ is dependent on HTT activity. Depletion of HTT also affected homologous recombination (HR)-mediated DSB repair but to a lesser (∼30%) extent (Supplementary figure 3D), suggesting that HTT depletion predominantly affects NHEJ-mediated DSB repair mechanism. Furthermore, increased phosphorylation of the DNA damage-response kinase, ATM, and of 53BP1 was observed in HTT-KD cells (Supplementary figures 3E and 3F), suggesting that persistence of DSBs in genome leads to aberrant activation of pro-degenerative DNA damage-response. By various measures, the HTT-KD cells consistently show significantly greater cell toxicities (Supplementary figures 3G and 3H). These combined observations suggest that HTT plays a key role in maintaining neuronal genome integrity—and neuronal health and viability—by facilitating DSB repair.

### BRG1-HTT facilitates NHEJ component recruitment at DSB sites

To test whether HTT-deficiency would impair DSB site recruitment of the HTT-associated NHEJ factors, we induced a DSB at a defined locus in chromosome 17 in control and HTT-deficient (HTT-KD) SH-SY5Y cells. We used liposome-mediated delivery of a CRISPR-CAS9 gRNA targeting a specific sequence on chromosome 17 to induce a DSB at that site^49^. After inducing a sequence-specific lesion, we followed the relative recruitments of key NHEJ factors at that specific DSB site in the HTT-KD and control cells using ChIP. Delivery of the sequence-specific CRISPR- CAS9 gRNA rapidly induced the site-specific DSBs in a time-dependent fashion as confirmed by a PCR-based ligation assay (Figure 5A). Further, immunostaining the SH-SY5Y cells treated with liposomes carrying CRISPR-CAS9 gRNA with γH2AX Ab also showed γH2AX-positive nuclear foci but such foci were not observed in cells treated with control empty liposome (Figure 5B; arrows), confirming that liposome-mediated delivery of CRISPR-CAS9 gRNA could introduce DSBs at specific genome loci. Subsequent ChIP analyses showed rapid recruitment of key NHEJ factors (53BP1, Ku70, Ku80, PNKP and DNA ligase IV) to the DSB in control SH-SY5Y cells, whereas recruitment of these NHEJ factors at the same DSB site in the HTT-KD cells was substantially reduced (Figure 5C), suggesting that HTT plays a dominant role in recruiting the NHEJ factors at DSB sites.

**Figure 5.**
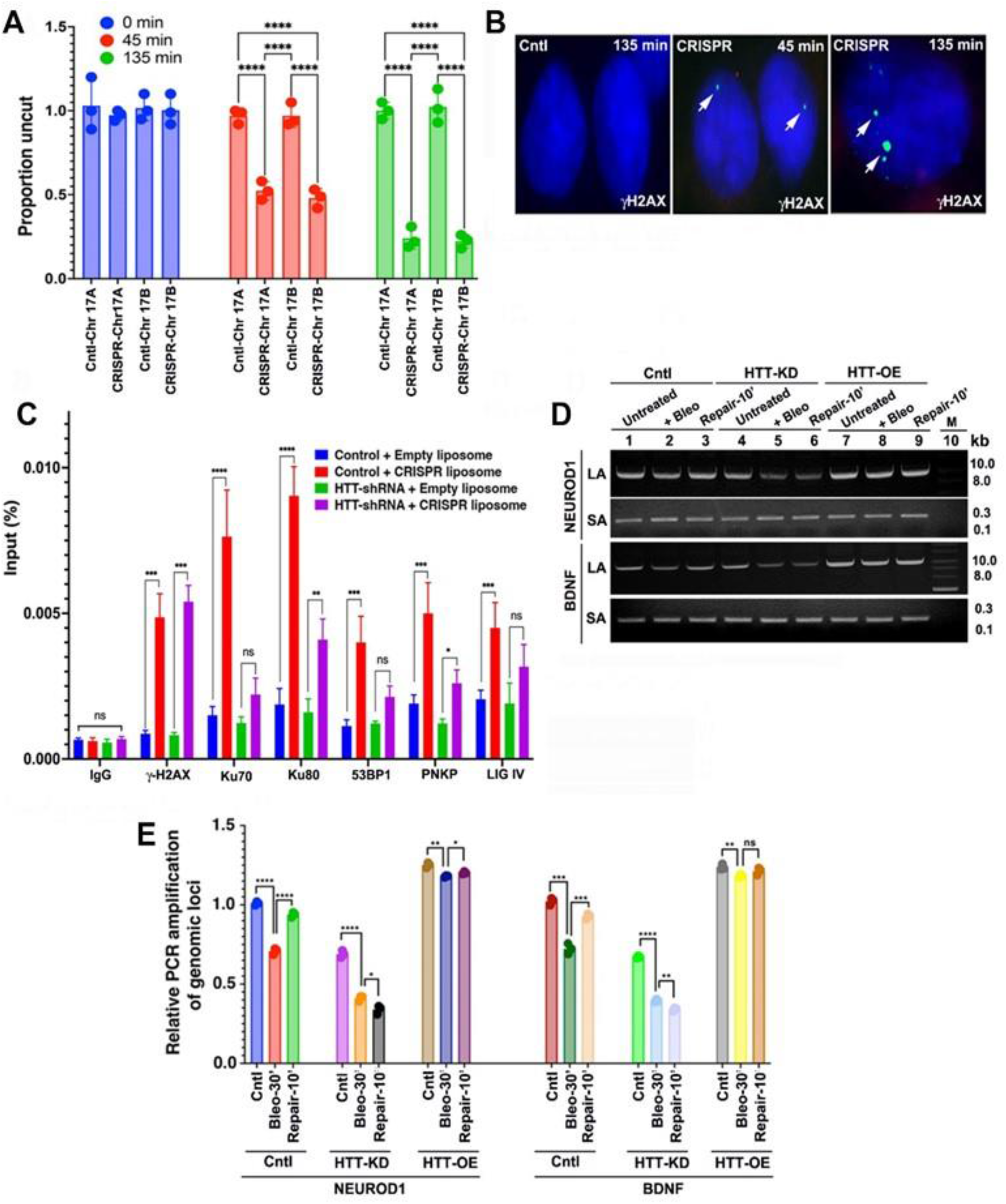
HTT facilitates recruitment of NHEJ factors at DSB sites. **(A)** Ligation-mediated PCR analysis at indicated time points after the addition of CRISPR- CAS9 gRNA containing liposomes measured the proportion of uncut DNA at two different sites of chromosome 17 (Chromosome 17: Chr17A and Chromosome 17B; Chr17B). (Mean ± SD. ****p<0.0001). **(B)** SH-SY5Y cells treated with either empty liposomes (Left panel) or liposomes carrying CRISPR-CAS9 gRNA (Central and right panels) for 45 or 135 minutes and cells analyzed by immunostaining with anti-γ-H2AX Ab. The γ-H2AX-positive nuclear foci shown by arrows. Nuclei stained with DAPI. **(C)** Low recruitment of NHEJ proteins in HTT-KD SH-SY5Y cells. DSBs were induced at specific locus of chromosome 17 (Chr17A) of control SH-SY5Y and HTT-KD SH- SY5Y cells using liposome-mediated delivery of CRISPR-CAS9 gRNA. Cells were harvested 135 minutes after adding liposomes, and ChIP performed to assess the relative recruitment of NHEJ proteins e.g., Ku70, Ku80, 53BP1, PNKP and DNA ligase IV at the DSB site in control and HTT-KD SH-SY5Y cells. Data represent Mean ± SD. **p<0.005; ***p<0.001; ****p<0.0001; and ns denotes not significant. **(D)** HTT depletion reduces DSB repair response to bleomycin. Control SH-SY5Y cells (Cntl; lanes 1 to 3), SH-SY5Y cells expressing HTT-RNAi (HTT-KD cells; lanes 4 to 6) and expressing the full-length human wtHTT-Q19 cDNA (HTT-OE cells; lanes 7 to 9) treated with bleomycin (BL; 5µg/mL; lanes 2, 3, 5, 6, 8 and 9) for 30 minutes. Cells were introduced with fresh medium to allow repair of the DSBs for 10’, cells harvested after 10 minutes of repair (R-10’; lanes 3, 6 and 9), DNA were isolated from each cell pellet, and various genomic DNA loci (10 kb or 0.2 kb genome segments encompassing NEUROD1 and BDNF genes) PCR amplified, and PCR products analyzed on agarose gels to determine relative PCR amplification efficacies. LA and SA denote long and short amplicon respectively. Lane 10: 1-kb DNA ladder. **(E)** HTT expression level correlates DSB repair. Relative levels of DNA damage in control SH-SY5Y cells (Cntl), HTT-KD cells, and HTT-OE cells, after treating the cells with BL (300 ng/mL) for 30 minutes, and 10 minutes after BL removal from the culture medium for allowing repair of the lesions (Recovery-10’); data represent means ± SD, *p<0.05; **p<0.005; ***p<0.001; ****p<0.0001; NS denotes not significant.

Induction of DSBs in HTT-KD or HTT-OE cells through treatment with bleomycin (BL; 5µg/mL, for 30 minutes), was used to evaluate DNA repair efficiency as a function of HTT expression. Cells treated with BL for 30 minutes were subsequently incubated in normal BL-free medium for 10 minutes to allow repair of the induced damage prior to rapid genomic DNA isolation. The genomic DNA was subjected to LA-QPCR analysis which revealed that HTT-deficient (HTT-KD) cells not only accumulated more lesions following BL exposure, but also poorly resolved lesions compared with control cells (Figures 5D; lanes 4-6 vs. lanes 1-3, and 5E). By contrast, HTT-OE cells exhibited greater resistance to DSB accumulation in response to BL insult and they quickly and completely repaired BL-induced lesions under identical experimental conditions (Figure 5D; lanes 7-9 vs. Lanes 1-3, and 5E). The delayed DSB repair and association of HTT with NHEJ proteins further support the notion that wildtype HTT protects neurons from genotoxic insults by promoting efficient DSB repair.

Previous studies showed that depleting BRG1 levels in epithelial U2OS cells induces DNA damage^27^, suggesting that BRG1 function is important in DNA repair mechanism. We therefore evaluated whether BRG1 similarly contributes to efficient NHEJ-mediated DSB repair and genome integrity in neuronal cells. We generated SH-SY5Y cells expressing either a BRG1 cDNA [BRG1-OE cells] or expressing a BRG1-shRNA [BRG1 knocked down; BRG1-KD cells]; BRG1 depletion or overexpression did not alter expression levels of key NHEJ proteins (Supplementary figure 4A). Immunostaining the cells with an anti-53BP1 Ab identified a widespread prevalence of 53BP1-positive nuclear foci in BRG1-KD cells (Supplementary figure 4B; upper panel; arrows) that was absent in control cells (Supplementary figure 4B; lower panel), suggesting that BRG1 depletion correlates with diminished DSB repair, resulting in persistence of DSBs. Consistently, LA-QPCR analyses of genomic DNA from the BRG1-KD cells revealed less efficient PCR amplification of various genome segments compared with controls (Supplementary figures 4C; lanes 4-6 vs. lanes 1-3, and figure 4D). As predicted, more efficient PCR amplification of the same genome segments was observed in BRG1-OE cells compared with controls (Supplementary figures 4C; lanes 7-9 vs. lanes 1-3, and figure 4D), reinforcing the notion that BRG1 acts in DSB repair essential for maintenance of genome integrity.

We next determined whether BRG1 facilitates recruitment of HTT at the DSB sites in the transcriptionally active genome. To test this idea, we simultaneously introduced two DSBs; one in a transcriptionally active chromosomal locus (locus A) and one in a transcriptionally inactive locus (locus B) in chromosome 1 in BRG1-deficient (BRG1-KD) and control SH-SY5Y cells using CRISPR-CAS9 gRNA as described above. We then followed recruitment of HTT at the DSB sites by using ChIP analysis. ChIP analyses revealed rapid and predominant recruitment of HTT at the DSB site located in the transcriptionally active locus (locus 1A) over the transcriptionally inactive locus in control cells (Supplementary figure 4E). In contrast, dramatically diminished HTT recruitment was observed at the same transcriptionally active chromosomal locus (locus 1A) in BRG1-depleted cells (Supplementary figure 4E). HTT recruitment at the transcriptionally inactive locus (locus 1B) did not change in BRG1-KD cells as compared with control cells (Supplementary figure 4E). To test whether HTT reciprocally helps in recruiting BRG1 at damage sites, we induced DSB at the same chromosomal locations in HTT-KD and control cells and assessed BRG1 recruitment at the DSB site in HTT-KD and control cells. The ChIP analysis revealed rapid and equally efficient recruitment of BRG1 at the DSB site in control cells, as well as in HTT-KD cells (Supplementary figure 4F). These observations indicate that BRG1 facilitates efficient recruitment of HTT at the DSB sites located in the transcriptionally active genome, and BRG1 deficiency impairs DSB site recruitment of HTT in the neuronal genome.

### Neuronal TC-NHEJ efficiently resolves DSBs in the transcriptionally active genome

A translocating transcription-coupled Repair (TCR) complex may sense lesions in the template DNA during transcription to initiate repair of the template DNA^20^. We therefore reasoned that if HTT is a functional component of TC-NHEJ, then HTT should be preferentially located within transcriptionally active (euchromatin) regions and recruited to DNA lesion sites. We therefore followed the relative occupancy/recruitment of HTT at the DSB sites at transcriptionally active locus A and transcriptionally inactive locus B regions described above before and after induction of the DSBs using ChIP analysis. ChIP analysis revealed a proportionally higher association of HTT with the DSB at the transcriptionally active genomic regions as compared with DSB locus within the transcriptionally inactive regions enriched for coding sequences—on both chromosomes—under stress-free condition (Figure 6A). When DSBs were simultaneously introduced at the chromosomal loci, HTT occupancy at the DSB sites located within the transcriptionally active regions (locus A) on either chromosome was increased over 2-fold, while recruitment of HTT to the DSB sites located within the transcriptionally silent heterochromatin regions of either chromosome (locus B) remained unaltered (Figure 6A).

**Figure 6.**
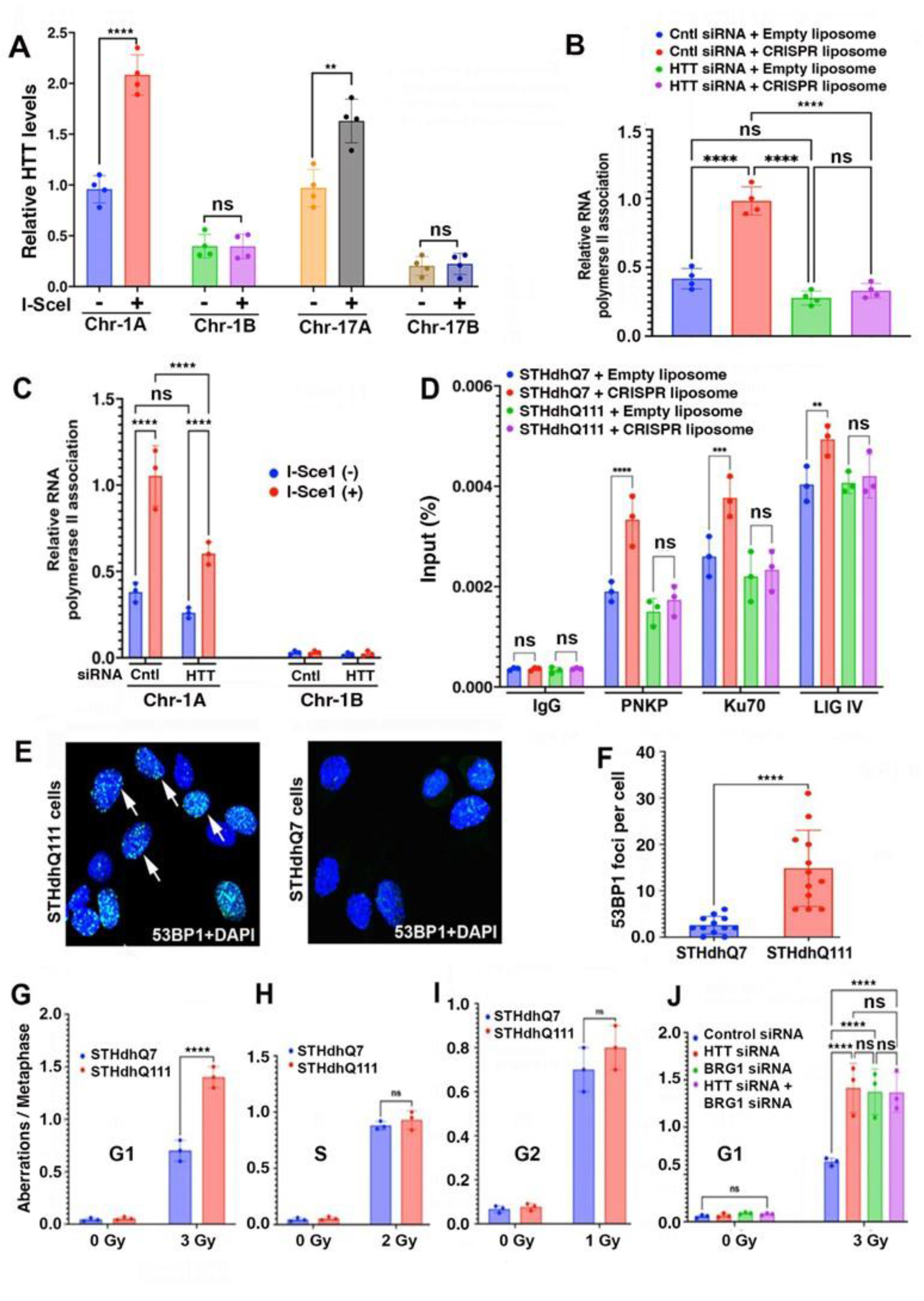
mHTT-mediated depletion of TC-NHEJ activity results in preferential accumulation of DSBs within the transcriptionally active genome *in vivo*. **(A)** Association of HTT with transcriptionally active DSB regions. DSBs were introduced within the transcriptionally active gene-rich locus (locus A) and in transcriptionally inactive gene-poor locus (locus B) in chromosome 1 and 17 (Chr-1 and Chr-17 respectively) in cells with and without induction of I-SceI restriction enzyme. Induced DSBs and relative occupancy/recruitment of HTT at the DSB sites within the transcriptionally active and in transcriptionally inactive chromosomal regions before and after inducing DSBs were determined by ChIP analysis. Data represent Mean ± SD. **p<0.005; ****p<0.0001; ns= not significant. **(B)** RNA polymerase II (POLR2A) levels at the DSB sites before and after CRISPR-CAS9 gRNA-mediated induction of DSB at Chr17A locus in SH-SY5Y cells with and without HTT-shRNA-mediated depletion of HTT. Mean ± SD. ***p<0.001. **(C)** RNA polymerase II (POLR2A) levels at the DSB sites before and after I-Scel induced DSBs at Chromosome 1 at transcriptionally active locus (Chr-1A) as well as in transcriptionally inactive locus (Chr-1B) were measured in cells with and without depletion of HTT. Data represent Mean ± SD. ***p<0.001. NS= not significant. **(D)** Robust recruitment of NHEJ factors in STHdhQ7 but not in mutant STHdhQ111 cells. STHdhQ7 and mutant STHdhQ111 cells were tested by liposome-mediated delivery of CRISPR-CAS12 gRNA, cells harvested 135 minutes after adding liposomes, and ChIP performed to assess relative recruitment of NHEJ proteins e.g., PNKP, Ku70, and DNA ligase IV at the DSB site in mutant STHdhQ111 and wildtype STHdhQ7 cells. Data represent Mean ± SD. **p<0.005; ***p<0.001; ****p<0.0001; ns= not significant. **(E)** DSBs persist in mutant STHdhQ111 cells but not in control STHdhQ7 cells. Mutant STHdhQ111 cells (left panel) and control STHdhQ7 cells (right panel) were analyzed by immunostaining with an anti-p-53BP1Ab (2675; Cell Signaling) to detect the presence of double strand breaks (DSBs) in nuclear genome. The p-53BP1-positive DSBs in genomic DNA (green puncta) within the nuclei (blue) of STHdhQ111 cells (left panel) are shown by arrows. Similar nuclear puncta not detected in control cell nuclei (right panel). Nuclei stained with DAPI (blue). **(F)** Relative amounts of 53BP1-positive puncta indicating DNA damage in mutant STHdhQ111 cells is high compared with control STHdhQ7 cells. Data represent means ± SD, ****p<0.0001. **(G)** Lower metaphase aberrations in wildtype STHdhQ7 compared to mutant STHdhQ111 cells. Wildtype STHdhQ7 and mutant STHdhQ111 cells in plateau phase were irradiated with 3 Gy, incubated for 18 hours post-irradiation, and G1-type aberrations examined at metaphase. Categories of asymmetric chromosome aberrations scored included dicentrics, centric rings, interstitial deletions-acentric rings, and terminal deletions. The frequency of chromosomal aberrations in STHdhQ111 cells after IR exposure were compared with control cells. Data represent means ± SD, ****p<0.0001). **(H)** Similar exponential phase aberrations in wildtype STHdhQ7 compared to mutant STHdhQ111 cells. Wildtype STHdhQ7 and mutant STHdhQ111 cells in exponential phase were exposed to 2 Gy IR, and metaphases harvested 3 hours post-irradiation and examined for chromosomal aberrations. The difference between chromatid and chromosomal aberrations induced by IR is not significantly higher in STHdhQ111 cells compared to control cells. ns= not significant. **(I)** Wildtype STHdhQ7 and mutant STHdhQ111 cells in exponential phase were irradiated with 1 Gy IR. Metaphases were harvested after 1-hour post-irradiation and analyzed for chromosomal aberrations. The differences in chromosomal aberrations between samples treated with IR are not statistically significant between STHdhQ111 and STHdhQ7 cells. Data represent means ± SD, **p<0.005. **(J)** Chromosome aberrations measured in control cells, HTT-depleted cells and HTT and BRG1 depleted cells before (0 Gy) and after irradiating the cells with IR (3 Gy). Data represent means ± SD, ****p<0.0001. ns = not significant.

As a positive control, we induced DSBs at the same transcriptionally active locus A in HTT-deficient (HTT-KD) cells and control cells and assessed the relative occupancy by RNA polymerase II (POLR2A), a key component of the TC-NHEJ complex before and after induction of the DSBs. The ChIP results reproduce the HTT findings where recruitment of POLR2A to the DSB site was substantially increased in control cells in response to DSBs, and recruitment of POLR2A at the DSB site is suppressed in HTT-deficient cells (Figure 6B). As an alternative approach, we introduced DSBs in transcriptionally active gene-rich Locus A and inactive genome locus B as before in HTT-deficient (HTT-KD) SH-SY5Y cells and control SH-SY5Y cells and followed the recruitment of POLR2A at these DSB sites. ChIP analyses show that the POLR2A is rapidly and efficiently recruited at the DSB site in control cells, but recruitment efficiency of POLR2A at the DSB sites is significantly impaired in HTT-KD cells. As expected, POLR2A recruitment at the DSB site in the transcriptionally inactive gene-poor locus was marginal (Figure 6C). These observations support a key role for HTT in recruiting and assembling a functional TC-NHEJ complex at the DSB site to facilitate TC-NHEJ-mediated DSB repair in neurons. Together, these data provide compelling evidence that HTT activity is central in the efficient recruitment of NHEJ factors at DSB sites within the transcriptionally active chromatin.

### mHTT-TC-NHEJ complex impairs DSB repair and increases DSB accumulation

To determine whether the presence of mHTT with a polyQ expansion interferes with recruitment of NHEJ factors at the DSB site, we induced DSBs at specific chromosomal loci in STHdhQ111 striatal cell model HD expressing mHTT and control STHdhQ7 cells expressing wtHTT^50^ and followed the recruitment of key NHEJ factors at the DSB sites. We introduced DSBs at a specific transcriptionally active genomic locus by using liposome-mediated delivery of a sequence-specific CRISPR-CAS12 gRNA in mutant STHdhQ111 or control STHdhQ7 cells and followed the recruitment of key NHEJ factors at the DSB site using ChIP analysis. The ChIP analysis showed efficient and rapid recruitment of key NHEJ factors, e.g., PNKP, Ku70 and DNA ligase IV (LIG IV) at the DSB site in control STHdhQ7 cells, and in contrast, recruitment of the same NHEJ factors at the DSB site in mutant STHdhQ111 cells is markedly impaired (Figure 6D). Consistently, immunostaining the cells with an anti-53BP1 Ab revealed widespread 53BP1- positive nuclear puncta in STHdhQ111 cells but not in control STHdhQ7 cells (Figures 6E and 6F). These observations strengthened the interpretation that mHTT in the TC-NHEJ complex inhibits efficient recruitment of the key NHEJ factors at the DSB sites impairing NHEJ, ultimately resulting in persistence of DSBs in mutant STHdhQ111 cells. Moreover, when exposed to IR the mutant STHdhQ111 cells showed substantially (∼2-fold) increased chromosome aberrations (Figures 6G-6I), suggesting intrinsically higher genomic instability in the mutant cells. To test whether reduction of both BRG1 and HTT further enhances chromosome aberrations, we simultaneously depleted both HTT and BRG1 in human cells (SH-SY5Y cells) and assessed chromosome aberrations at metaphase. Chromosome analyses revealed that combinatorial depletion of HTT and BRG1 in cells did not further enhance chromosome aberrations compared with the chromosome aberration that was observed by depleting HTT alone (Figure 6J). These observations support the idea that HTT is epistatic to BRG1 and mechanistically function in the same pathway in DSB repair as well as genome integrity.

We next used zQ175 knock-in mouse brain tissue expressing an expanded repeat within the endogenous mouse locus^32^ to determine whether mHTT in the TC-NHEJ complex impairs DSB repair *in vivo*, and whether DSBs accumulate at early stages of disease progression before the onset of overt motor dysfunction. We therefore analyzed pre-motor symptomatic (12-week-old) zQ175 heterozygous and age-matched control mouse brain sections (striatum) for the presence of DSBs in neuronal genome by immunostaining the brain sections with an anti-53BP1 antibody. A widespread distribution of 53BP1-positive nuclear foci was observed in zQ175 brain sections (Supplementary figure 5A; arrows) but not in control (Supplementary figure 5B), suggesting that spontaneous DSBs in the neuronal genome associated with routine neuronal activities remain largely unresolved likely due to impaired NHEJ in HD.

If TC-NHEJ becomes dysfunctional at early stages of disease progression, we should expect that DNA repair deficiency and DNA damage accumulation should occur early during the prodromal stages of disease progression. To test this hypothesis, we isolated genomic DNA from the presymptomatic 12-week-old zQ175 transgenic^32^ and age-matched control mouse cortex. We then quantified the relative DNA strand breakage in actively transcribed versus non-transcribed genomic regions. The LA-QPCR analyses revealed about 40-50% lower PCR amplification efficacy of genomic segments that harbor actively transcribed genes regulating synaptic function, vesicular transport, and calcium homeostasis in asymptomatic zQ175 cortical tissue compared to controls (Supplementary figures 5C and 5D). The evidence for impaired TC-NHEJ activity in 12- week-old mice manifesting as persistent DSBs *in vivo* may represent early stages of disease. Furthermore, the marginal difference in PCR amplification efficacies detected by LA-QPCR analysis targeting transcriptionally inactive chromatin (e.g., MyoD1 and MyoG1) in zQ175 cortical tissue compared with controls, indicates that DNA damage might be lesser, or the DNA repair process is generally less robust in transcriptionally active gene-rich genome compared with transcriptionally inactive gene-poor genome regions (Supplementary figures 5E and 5F). These findings suggest that the HTT-containing TC-NHEJ complex repairs DSBs in the transcriptionally active genome, while mHTT-mediated inactivation of the TC-NHEJ complex biases the distribution of persistent lesions to transcriptionally active “gene-rich” genomic areas.

Immunostaining of both iPSC-derived HD and control neurons^51^, with an anti-53BP1 Ab showed a widespread 53BP1-positive nuclear foci in HD but not in control iPSC-derived neurons (Supplementary figures 6A and 6B; arrows). This suggested DSBs are not efficiently resolved and persist in iPSC-derived HD primary neurons. We therefore analyzed HD postmortem brain sections (n=6) and control non-diseased postmortem brain (putamen) sections by immunostaining the brain sections with anti-53BP1 antibody (a marker for DSBs) to determine whether DSBs remain unrepaired and persist in HD brain. Immunostaining analyses revealed widespread presence of 53BP1-positive nuclear foci in brain sections from HD patients (52-year-old; Grade 2- and 56-year-old; grade 3), but not in age-matched normal (non-diseased) controls (Supplementary figures 6C and 6D; arrows), reflective of an impaired DSB repair mechanism in the HD patients’ brain. The LA-QPCR analysis of the genomic DNA isolated from the postmortem HD caudate-region brain tissue showed significantly reduced amplification products for various genome segments, consistent with broadly distributed persistent DSBs in HD brains (Supplementary figures 6E and 6F). These data suggest that mHTT’s interference with NHEJ represents a consistent mechanism across species, responsible for persistent accumulations of DSBs in HD patient-derived primary neurons and HD patient brain tissue.

### wtHTT expression improves genome integrity, neurotoxicity, motor defects, and lifespan in HD cell and *Drosophila* models

We tested whether wtHTT expression in a cell model of HD stimulates DNA repair by stoichiometrically increasing the number of active TC-NHEJ complex recruited to chromatin in the mutant cells. We assessed DNA lesions in mutant STHdhQ111 cells, a characterized cell model of HD, and control STHdhQ7 cells^50^. The LA-QPCR analysis of the genomic DNA from mutant STHdhQ111 cells showed significantly reduced PCR amplification products targeting several genome segments in the mutant cells compared with control cells, suggesting higher incidence of DNA lesions in the mutant cells (Figures 7A; lanes 4-6 vs. lanes 1-3; and 7B). We next expressed wtHTT-Q24 (wildtype full-length HTT encoding 24 glutamines) into the mutant STHdhQ111 cell background, isolated genomic DNA and conducted LA-QPCR analysis on the genomic DNA which revealed enhanced PCR amplification of the same genome segments (Figures 7A; lanes 7- 9 vs. lanes 4-6; and 7B). The findings suggest that increasing wtHTT expression in the mutant background can compensate for the faulty HTT and improve overall genome integrity.

**Figure 7.**
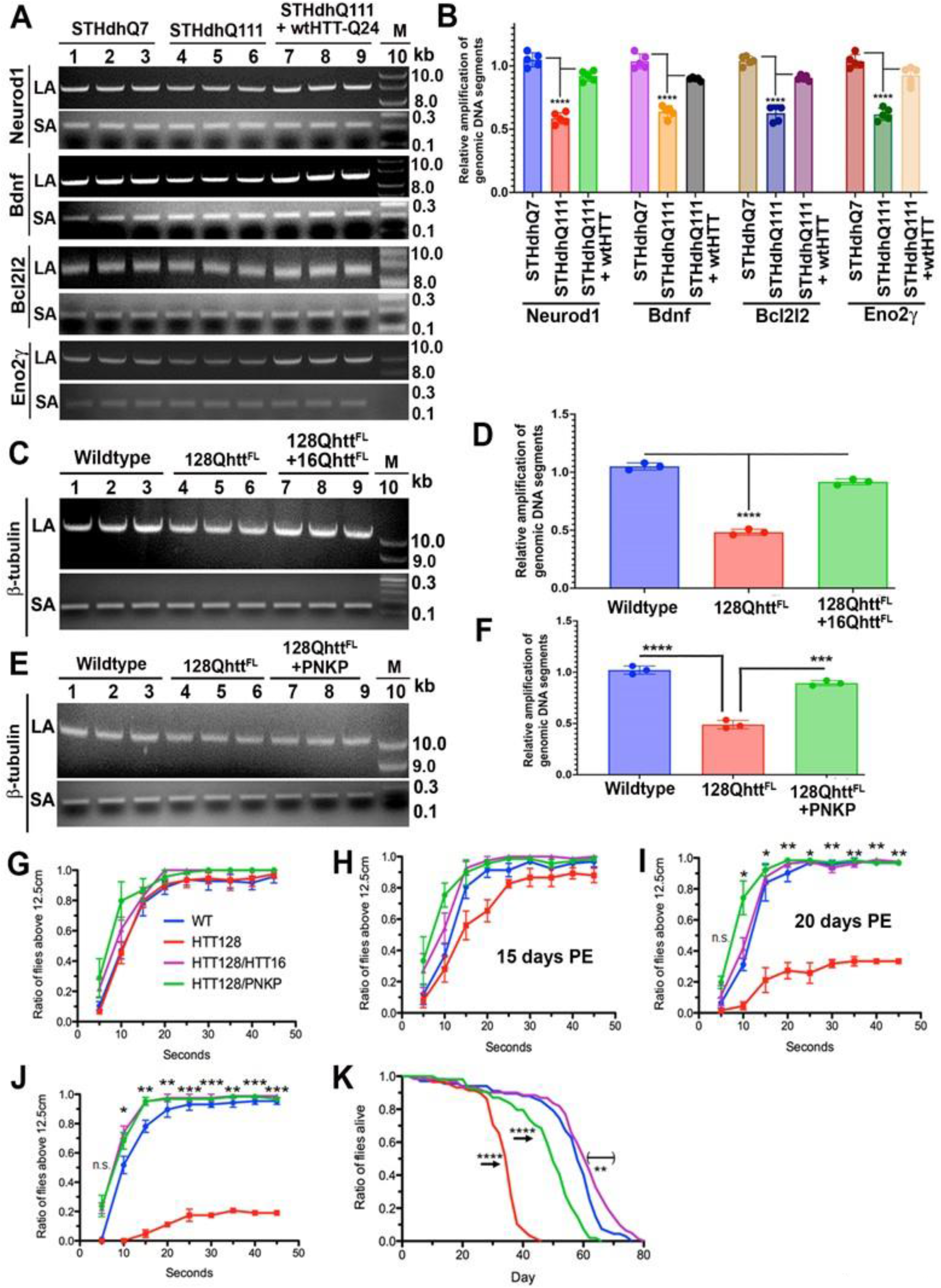
Increasing wtHTT expression in HD cell and *Drosophila* models reduces DNA damage while also improving genome integrity, motor defects and longevity, and neuronal gene expression. **(A)** Genomic DNA isolated from the control STHdhQ7 (lanes 1 to 3), mutant STHdhQ111 cells (lanes 4 to 6) or mutant STHdhQ111 cells expressing full-length wildtype human HTT carrying 24 glutamines (wtHTT-Q24; lanes 7 to 9), and various genomic DNA segments encompassing the Neurod1, Bdnf, Bcl2l2 and Eno2α genes PCR amplified; and PCR products analyzed on agarose gels, and intensity of the DNA bands quantified. LA denotes long amplicon (8 to 10 kb), SA denotes short amplicon (0.2 to 0.3 kb). Lane 10: 1-kb ladder. **(B)** Relative PCR amplification of various genomic DNA segments from the Neurod1, Bdnf, Bcl2l2 and Eno2α gene loci in STHdhQ7, STHdhQ111, and STHdhQ111 cells ectopically expressing full-length human wtHTT-Q24. Six technical replicates are used for the calculations. Data represent mean ± SD, ****p<0.0001. **(C)** Genomic DNA isolated from wildtype flies, mutant UAS-128Qhtt^FL^, and UAS- 128Qhtt^FL^ *drosophila* flies expressing human wildtype full-length HTT-Q16 and the β- tubulin locus (∼10 kb) PCR amplified, and PCR products analyzed on agarose gel and the intensity of the DNA bands quantified. Twenty-five biological replicates and three technical replicates were used for this measurement. Data represent mean ± SD, ****p<0.001 (UAS-128Qhtt^FL^ vs. wildtype) and (mutant flies expressing wildtype full-length human HTT (wtHTT-Q16) vs. mutant HD flies). **(D)** Relative PCR amplification of β-tubulin loci in control wildtype (WT) flies, mutant UAS-128Qhtt^FL^, and mutant UAS-128Qhtt^FL^ d*rosophila* ectopically expressing wtHTT-Q16. Twenty-five biological replicates and three technical replicates were used for this measurement. Data represent mean ± SD, ****p<0.0001 (UAS-128Qhtt^FL^ vs. WT) and (mutant flies expressing exogenous wtHTT vs. mutant flies). **(E)** Genomic DNA isolated from wildtype fly heads (lanes 1 to 3), HD transgenic UAS- 128Qhtt^FL^ fly heads (lanes 4 to 6) and the UAS-128Qhtt^FL^ fly heads expressing human full-length PNKP (lanes 7 to 9), and DNA damage assessed by LA-QPCR. β-tubulin genomic DNA locus of *drosophila* genome PCR amplified, and PCR products analyzed on agarose gels and quantified. LA denotes long amplicon and SA denotes short amplicon. Lane 10: 1-kb DNA ladder. **(F)** Relative PCR amplification of β-tubulin loci in control wildtype (WT) flies, mutant UAS-128Qhtt^FL^, and mutant UAS-128Qhtt^FL^ *drosophila* ectopically expressing human PNKP. Twenty-five biological replicates and three technical replicates were used for this measurement. Data represents mean ± SD, ****p<0.0001 (UAS-128Qhtt^FL^ vs. WT) and (mutant flies expressing exogenous PNKP vs. mutant flies). **(G to J)** Climbing properties of *Drosophila* that reflects the motor coordination measured after (**G**) 10 days, (**H**) 15 days, (**I**) 20 days, and (**J**) 25 days post eclosion in wildtype (WT), UAS-128Qhtt^FL^ and UAS-128Qhtt^FL^ flies either expressing full-length human wildtype (WT) HTT-Q16 (wtHTT-Q16) or human PNKP measured by standard climbing properties of flies. Relative motor function of wildtype, UAS-128Qhtt^FL^, and UAS-128Qhtt^FL^ *drosophila* expressing full-length wtHTT-Q16 or PNKP are shown. *p < 0.05, **p < 0.01, ***p < 0.001, ****p < 0.0001 (UAS-128Qhtt^FL^ vs. WT) (note that these significance values were calculated solely from the repeated experimental trials and that an average of 25 flies were used in each trial for each group at each time point). **(K)** Relative lifespans of *Drosophila* measured in wildtype, UAS-128Qhtt^FL^ and UAS- 128Qhtt^FL^ flies ectopically expressing either human PNKP or wildtype full-length HTT-Q16. Relative longevities of the wildtype, UAS-128Qhtt^FL^, and UAS-128Qhtt^FL^ *drosophila* expressing either exogenous PNKP or full-length human wtHTT-Q16 are shown. **p < 0.01, ****p < 0.0001 (note that start numbers of same day eclosion adult *Drosophila* for each group were 67 for WT, 58 for HTT128, 61 for HTT128/HTT, and 54 for HTT128/PNKP).

To determine if increasing wtHTT expression *in vivo* would stimulate DNA repair, improve genome integrity, and lessen neurotoxicity, we used a *Drosophila* model of HD that recapitulates many HD-like phenotypes^52, 53^. First, we expressed mHTT (128Qhtt^FL^) or wtHTT (16Qhtt^FL^) in *Drosophila* neurons using the UAS-Gal4 system^54^. The BG-380 gal4 driver^55^ was used to express UAS-16Qhtt^FL^ in the neurons of the 128Qhtt^FL^-expressing flies. Genomic DNA isolated from fly heads was analyzed for DNA damage. As anticipated, LA-QPCR assays revealed less efficient PCR amplification of a genomic DNA segment isolated from the mutant flies (20-day old) compared to age-matched wildtype flies (Figure 7C; lanes 4-6 vs. lanes 1-3, and 7D), reflective of increased accumulated genome damage in mutant flies. However, increasing wtHTT expression in the mHTT background (UAS-128Qhtt^FL^) dramatically reduced the level of DNA damage (Figures 7C; lanes 7-9 vs. lanes 4-6, and 7D) supporting the hypothesis that wtHTT promotes DNA repair and genome integrity. HTT DNA repair functions are therefore a logical potential underlying mechanism for wtHTT in conferring neuroprotection *in vivo*^40, 41, 44, 47^.

wtHTT regulates many cellular processes including endocytosis, and transport of organelles and vesicles in the axons and these processes disrupted by mHTT^56, 57^. We therefore tested whether stimulating DNA repair per se can improve genome integrity and motor defects using an *in vivo* model. To do this, we generated a transgenic fly line expressing human PNKP under the control of the Gal4 promoter (BG-380-gal4) in the mHTT fly background. As predicted, mutant flies showed greater DNA damage (Figures 7E; lanes 4-6 vs. lanes 1-3, and 7F). Importantly, introduction of the transgenic PNKP expressed in neurons of the mutant fly (UAS- 128Qhtt^FL^) significantly reversed the level of genome damage (Figures 7E; lanes 7-9 vs. lanes 4- 5 and 7F). These results support and extend the cell culture findings that increased DNA damage is likely due to ineffective DNA repair. Conversely, neuronal genome integrity could be recovered in the *in vitro* as well as *in vivo* model of HD through harmonizing the TC-NHEJ complex with wtHTT and/or PNKP.

We then harnessed the fly model to determine whether the proposed role for wtHTT in fixing DNA lesions to maintain genome integrity would also reverse the motor defects and increase longevity of the mutant flies. Increasing wtHTT or PNKP expression in mutant flies prevented onset of motor defects (Figures 7G to 7J) and substantially increased their lifespan (Figure 7K). Moreover, enhancing wtHTT or PNKP expression in mutant flies largely normalized expression of many genes whose expression is altered in the mutant flies. This included genes with human homologues known to be involved in neurodegenerative diseases, such as β-secretase (BACE1) (Supplementary figures 7A and 7B). These collective data show that wtHTT plays a key role in organizing a functional NHEJ DSB repair complex in neurons whereas polyQ expansion compromises this repair complex degrading genome integrity, transcription, neuronal survival, and motor function in HD.

## DISCUSSION

HTT is a large multifunctional protein, so how may its function in TC-NHEJ inform and advance our knowledge of mHTT impacts and HD research? The postmitotic status of most neurons coupled with their high metabolic activities renders their genomes exquisitely susceptible to DNA damage by endogenous genotoxic metabolites^1, 7^. DNA breaks, genome instability, and transcription stress that are associated with oxidative metabolism are linked to widespread functional decline and segmental aging^58^. Increasing evidence suggests that impaired or faulty DNA repair undermines genome integrity with concomitant aberrant gene expression, neuronal dysfunction, and premature death of neurons^3, 4, 7, 59^. Oxidized DNA bases, SSBs, and DSBs are repaired by the BER pathway in the CNS^60^. Nucleotide excision repair (NER), mismatch repair (MMR), and NHEJ pathways are also important in resolving DNA lesions in neurons^4, 7, 61^. In mammalian cells, DSBs are repaired by NHEJ and homologous recombination (HR)^62, 63^ with NHEJ occurring throughout the cell cycle except for M-phase. Consequently, NHEJ is the major DSB repair mechanism in neurons^62^.

Importantly, the wide-spread occurrence of DSBs in the brains of patients is associated with multiple neurodegenerative disorders, such as Alzheimer’s disease^64^, Parkinson’s disease^65, 66, 67^, Frontotemporal lobar degeneration/Amyotrophic Lateral Sclerosis (ALS)^68, 69^, or Spinocerebellar ataxia types 3 and 7 ^70, 71, 72^. Mice expressing a kinase-dead DNA-PKcs^73^, and inactivation of genes encoding NHEJ proteins e.g., DNA-PKcs^74^, LIG IV^75, 76^ or XRCC4^76^ are embryonically lethal with neuronal apoptosis suggesting that the NHEJ pathway is critically important for health and survival of adult neurons. Thus, there is evidence that inefficient NHEJ may contributing to these pathologies. In fact, either diminished levels or activities of critical NHEJ factors occur in Alzheimer’s disease^77, 78^ and ALS^68^. Consistently, our results show that mHTT disrupts the functional integrity of a TC-NHEJ complex in neurons with repercussions on DNA repair and genome integrity critical for maintaining normal neuronal function.

### HTT-BRG1 scaffolds an efficient TC-NHEJ complex in neurons

Neurons have extreme vulnerability to transcription-associated DSBs due to their high metabolic activity coupled to high transcriptional activity with over two-thirds of genes transcribed and much of the genome in an open damage susceptible chromatin state. Yet, neurons lack homologous recombination (HR)^5^, potentially predisposing them to accumulating DSB lesions with age. Our results indicate that efficient recruitment of HTT and key NHEJ factors at the DSB sites in the gene-rich transcriptionally active genome is BRG1-dependent. Furthermore, we determined that BRG1-deficiency adversely impacts recruitment of HTT and NHEJ proteins at DSB sites. Notably, suppressing and overexpressing either HTT or BRG1 in neuronal cells increases and decreases DSBs, respectively, suggesting why wtHTT acts as a strong neuroprotective factor against the metabolically derived genotoxic insults^40, 41, 44, 46, 47^. Consistent with HTT scaffolding a TC-NHEJ complex, we find that increasing wtHTT expression leads to increased assembly and functionality of the TC-NHEJ complex. Importantly, this complex restores TCR to improve genome integrity plus expression of neuronal genes, decrease motor defects, and increase lifespan in an established *Drosophila* HD model. In contrast, mHTT disrupts TC-NHEJ functionality. This repair defect results in accumulation of DSBs, rendering neurons more vulnerable to stress-mediated damage and premature loss. These results support a model where the translocating TC-NHEJ complex pauses at the DSB sites in normal healthy neurons and facilitates phosphorylation and subsequent recruitment of the NHEJ factors in a HTT-facilitated process at the DSB sites to maintain genome integrity (as depicted in Supplementary figure 7C).

### Impacts of mHTT PolyQ expansion on BRG1 activity and chromatin dynamics

BRG1 and BRG1-associated factors (BAFs) catalyze ATP-dependent reactions promoting histone assembly/disassembly from DNA altering chromatin architecture to modulate transcription during neuronal development^24, 25, 79^. As HTT interacts with BRG1 to scaffold the TC-NHEJ complex, identification of BAFs present in the neuronal HTT-BRG1 complex may better inform how this complex affects chromatin dynamics in transcription-coupled DNA repair, and how polyQ expansion in mHTT interferes with chromatin structure and transcription in HD. Multiple specific mechanisms can now be investigated in future studies. Expanded polyQ sequences in mHTT may degrade the HTT-BRG1 functional complex, leading to erosion of the neuronal transcriptional repertoire and subsequent HD. For example, mHTT may repress transcription by directly interfering with the activity of neuron-specific transcription factors^80, 81, 82^. mHTT may disrupt chromatin dynamics by altering histone post-translational modifications known to modulate gene expression, including inappropriate changes in histone acetylation by histone acetyltransferases (HATs), resulting in transcriptional dysregulation in HD^83, 84^. Similarly, the failure of timely clearance for various transcription factors due to impaired proteostasis might also contribute to transcriptional dysregulation in HD^84, 85, 86^.

### HTT scaffolds a TC-NHEJ complex to maintain the transcriptionally active genome

Our collective multi-disciplinary findings indicate that a neuronal HTT scaffolded TC-NHEJ complex acts predominantly in the transcriptionally active gene-rich genome and less so the non-transcribed regions. Thus, impairment of this complex by polyQ expansion in mHTT disrupts both DNA repair and transcription to reduce neuronal health and function as a potential foundational mechanism for HD. Progressive accumulation of DSBs within gene coding regions, promoters, and regulatory elements of actively transcribed genes will cumulatively impair essential neuronal functions and ultimately erode overall neuronal health and function culminating in neurotoxicity and disease progression in HD. Our data and insights thus suggest that reduced DSB repair due to a combination of decreased HTT function and the presence of mHTT in the TC-NHEJ complex significantly contribute to neurotoxicity and neurodegeneration in HD. It will therefore now be important to test if DSB damage in adult neurons can be resolved by stimulating DSB repair and furthermore if strategies designed to stimulate repair before neurons reach the tipping point may aid prevention of symptomatic disease.

## LIMITATIONS OF THE STUDY

These results identify a functionally relevant TC-NHEJ complex scaffolded by HTT that is impaired by mHTT promoting unrepaired DSBs that degrade transcription and neuronal genome integrity consistent with HD progression. Molecular mechanisms for HTT recruitment, scaffolding, and impairment by mHTT remain to be determined. The AF3 structure prediction places HTT between BRG1 and PNKP and capped by Ku70/80, but this is a computational hypothesis to guide future experimental testing. Also, these data do not link HTT’s NHEJ role to a mechanism for CAG triplet repeat expansions. Furthermore, genetic association studies and other data suggest non-NHEJ DNA repair proteins, which may have crosstalk with DSB repair (including FAN1, alternative RPA, and MSH3), are linked to CAG triplet repeat expansion and modify age at onset for HD. Despite the interwoven complexity in related DNA repair processes that can evidently influence timing and progression for HD, the existing data imply that mechanisms for CAG repeat expansion may be separable from the mHTT pathogenesis caused by transcription-associated DSB repair defects in brain reported here. Dissecting potential interrelationships among DNA repair processes in HD onset and progression as well as possible implications for transcription coupled repair defects in other polyglutamine (polyQ) diseases require further investigations.

## ACKNOWLEDGEMENTS

This research was supported by National Institutes of Health grants RO1 NS130830 and R01 EY026089-01A1 to P.S.S.; Hereditary Disease Foundation (HDF) grant to P.S.S; R35 CA220430 to J.A.T.; R35 NS116872 and R01 NS090390 to L.M.T. and Hereditary Disease Foundation Fellowships to C.G. and D.F. P.; R56 NS105681, and Alzheimer’s Association grant to Y.P.W., National Institutes of Health grants R01ES033682 to C.E., Alzheimer’s Association grant AARF- 22-967275 to S.G., and RO1ES034542 to R.K.P. We acknowledge NeuroBiobank for the HD and control brain tissue samples. We thank Junjei Chen at MD Anderson Cancer Research Center for allowing us to use X-ray irradiator for irradiating cells.

## AUTHOR CONTRIBUTIONS

S.P., S.C., R.K.P., T.K.P. generated the ChIP, ChIP-re-ChIP, WB, IP, PLA, and LA-QPCR analyses. S.P., D.F.P., and P.S.S generated various plasmid constructs and cell models of HD, performed LA-QPCR, and qPCR analyses. S.P and S.G. performed the MS analysis. C.L.T and J.A.T did computational modeling and structural analyses and contributed to writing. N.Z., A.T., and L.N. performed cell toxicity measurements and protein interaction studies by PLA. D.F.P, C.S., and L.M.T. characterized and provided the iPSC-derived neurons, provided brain tissues from HD mouse models, and helped draft the manuscript. K.B and Y.P.W. generated the *Drosophila* expressing human PNKP and performed the motor function and longevity analysis and helped draft the manuscript. L.F., R.K.P. K.S.R. and T.K.P. produced the liposomes carrying the CRISPR-Cas9 gRNA and performed DNA damage analysis. S.Y. performed the confocal image analyses. Project leader P.S.S. wrote and edited the final manuscript with substantial input from C.E., K.S.R, J.A.T., T.K.P, and L.M.T.

## COMPETING INTERESTS

The authors declare there are no competing interests.

## STAR METHODS

### Plasmid construction

Plasmid carrying *PNKP* and *full-length wild-type HTT* cDNA were purchased from Origene, USA, and PNKP or HTT cDNA were PCR-amplified with appropriate primers and sub- cloned into plasmid pSELECT-Puro-mcs (InvivoGen, USA) with appropriate linkers to construct plasmids pRSG-PNKP and pRSG-wtHTT, expressing PNKP or HTT respectively. These plasmids were transfected into SH-SY5Y cells, and the stable clones were selected for puromycin (2 μg/ml) resistance, and expression of exogenous PNKP or HTT were verified by western blots. Plasmid encoding human DNA ligase IV was purchased from Origene (USA) and the ligase IV cDNA was PCR-amplified with appropriate primers and was subcloned into plasmid pCMV-Myc-N (Takara Bio). Plasmid pBacMam2-DiEx-LIC-C-flag-huntingtin full length_Q24 (Addgene plasmid # 111742; http://n2t.net/addgene : Addgene _111742), pBacMam2-DiEx-LIC-C- flag_huntingtin_full-length_Q19 (Addgene plasmid # 111741; http://n2t.net/addgene: 111741; RRID:Addgene_111741), pBacMam2-DiEx-LIC-C-Flag_huntingtin_full-length_Q79 (Addgene plasmid# 111729; http://n2t.net/addgene: 111729; RRID: Addgene_111729), pBacMam2-DiEx-LIC-C-flag_huntingtin_full-length_Q109, (Addgene plasmid # 111730; http://n2t.net/addgene: 111730; RRIDAddgene_111730, pBacMam2-DiEx-LIC-C-flag_huntingtin_full-length_Q109, (Addgene plasmid # 111730; http://n2t.net/addgene: 111730; RRID: Addgene_111730), and pBacMam2-DiEx-LIC-C-C-flag_huntingtin_full-length_Q145 (Addgene plasmid#111731; http://n2t.net/addgene: 111731; RRID: Addgene_111731 were gifts from Cheryl Arrowsmith. The full-length huntingtin cDNA was PCR-amplified from these plasmids, and the PCR products were sub-cloned into plasmid pcDNA3.1 (Invitrogen, USA) using appropriate DNA linkers to construct plasmids pRGS-wtHTT-Q19, pRGS-wtHTT-Q24, pRGD-mHTT-Q79, pRSG-mHTT-Q109 and pRGS-mHTT-Q145, expressing full-length FLAG-tagged HTT. Plasmid encoding human DNA ligase IV and XRCC1 cDNA were purchased from Origene, USA, and the cDNA sequences were PCR-amplified with appropriate primers, and the PCR products were sub-cloned into plasmid pCMV-Myc-N (Takara Bio, USA) to constructs recombinant plasmids expressing Myc-tagged proteins. Plasmid pCMV-PARP1-3xFlag-WT was a gift from Thomas Muir (Addgene plasmid # 111575; http://n2t.net/addgene: 111575; RRID:Addgene_111575); plasmid N-Myc-53BP1 WT pLPC-Puro was a gift from Titia de Lange (Addgene plasmid # 19836; http://n2t.net/addgene: 19836; RRID:Addgene_19836); plasmid pEGFP-C1-FLAG-Ku80 was a gift from Steve Jackson (Addgene plasmid # 46958; http://n2t.net/addgene: 46958; RRID:Addgene_46958); plasmid pEGFP-C1-FLAG-Ku70 was a gift from Steve Jackson (Addgene plasmid # 46957; http://n2t.net/addgene: 46957; RRID:Addgene_46957); plasmid pcDNA3.1(+)Flag-His-ATM was a gift from Michael Kastan (Addgene plasmid # 31985; http://n2t.net/addgene: 31985; RRID:Addgene_31985). Plasmid pEGFP-C1-FLAG-XRCC4 was a gift from Steve Jackson (Addgene plasmid #46959; http://n2t.net/addgene:46959; RRID: Addgene_46959). The plasmid hu-DNA-PKcs was a gift from Katheryn Meek (Addgene plasmid# 83317). The plasmid pCMV5 BRG1-Flag was a gift from Joan Massague (Addgene plasmid # 19143; http://n2t.net/addgene: 19143; PRID: Addgene_19143).

The full-length cDNA sequences of Ku70, Ku80, XRCC4, DNA Ligase IV, ATM, and 53BP1 from these plasmids were PCR amplified with appropriate primers and sub-cloned into pCMV-Myc-N (Takara Bio, USA) to construct recombinant plasmids expressing Myc-tagged proteins. The full-length human BRG1 cDNA sequences from the plasmid pCMV5-BRG1-FLAG were PCR-amplified with appropriate primers and the PCR product was sub-cloned into pCMV-Myc-N plasmid (Takara Bio, USA) to construct recombinant plasmid pMyc-BRG1, expressing Myc-tagged BRG1. Various domains of BRG1 were PCR amplified with appropriate primers and subcloned into pCMV-Myc-N plasmid to construct recombinant plasmids that express various domains of BRG1. The sequence integrity of the clones was verified by sequencing, and expression of the proteins were checked by western blots with appropriate antibodies.

### Mass Spectrometry

Huntingtin (HTT) interacting proteins extracted from nuclear fractions were subjected to mass spectrometry analysis at the Mass Spectrometry Facility of the University of Texas Medical Branch, following the protocol described previously^87^. Briefly, HTT co-immunoprecipitated proteins were thoroughly washed using 50 mM Triethylammonium bicarbonate (TEAB), pH 7.1 buffer, then solubilized in a solution containing 5% SDS and 50 mM TEAB, pH 7.55. This mixture was incubated at room temperature for 30 minutes. The supernatants were then transferred to a fresh tube, reduced with 10 mM Tris (2-carboxyethyl) phosphine (TCEP) (Thermo Scientific, Cat # 77720) buffer, and incubated at 65°C for 10 minutes. Following cooling to room temperature, 3.75μL of 1M iodoacetamide was added, and the mixture was incubated in the dark for 20 minutes. The reaction was quenched with 0.5μL of 2M DTT. Subsequently, 5μL of 12% phosphoric acid was introduced to the 50μL protein solution, followed by the addition of 350μL of binding buffer (90% Methanol, 100 mM TEAB, pH 7.1). The mixture was then processed through an S-Trap spin column (Protifi, Farmingdale, NY) using a bench-top centrifuge (30 seconds at 4,000g). The spin column was washed three times with 400μL of binding buffer and centrifuged at 1200 rpm for 1 minute. The protein samples were digested with trypsin (Promega, #V5280, Madison, WI) at a 1:25 ratio in 50 mM TEAB, pH 8, and incubated at 37°C for 4 hours. Peptides were eluted sequentially with 80μL of 50 mM TEAB, 80μL of 0.2% formic acid, and 80μL of 50% acetonitrile with 0.2% formic acid. The pooled peptide solution was concentrated in a speed vacuum (room temperature, 1.5 hours) and reconstituted in 2% acetonitrile, 0.1% formic acid, and 97.9% water before being loaded into an autosampler vial.

Peptide mixtures were analyzed by nanoflow liquid chromatography-tandem mass spectrometry (nanoLC-MS/MS) using a nano-LC chromatography system (UltiMate 3000 RSLCnano, Dionex), coupled on-line to a Thermo Orbitrap Fusion mass spectrometer (Thermo Fisher Scientific, San Jose, CA) through a nanospray ion source (Thermo Scientific). A trap and elute method were used. The trap column was a C18 PepMap100 (300um X 5mm, 5um particle size) from Thermo Scientific The analytical columns was an Acclaim Pep Map 100 (75um X 25 cm) from (Thermo Scientific). After equilibrating the column in 98% solvent A (0.1% formic acid in water) and 2% solvent B (0.1% formic acid in acetonitrile (ACN)), the samples (2 µL in solvent A) were injected onto the trap column and subsequently eluted (400 nL/min) by gradient elution onto the C18 column as follows: isocratic at 2% B, 0-5 min; 2% to 24% B, 5-86 min; 24% to 44% B, 86-93 min; 44% to 90% B, 93-95 min; 90% B for 1 minute, 90% to 10% B, 96-98 min; 10% B for 1 minute 10% to 90% B, 99-102 min 90% to 4% B; 90% B for 3 minutes; 90% to 2%, 105-107 min; and isocratic at 2% B, till 120 min.

All LC-MS/MS data were acquired using XCalibur, version 2.5 (Thermo Fisher Scientific) in positive ion mode using a top speed data-dependent acquisition (DDA) method with a 3 second cycle time. The survey scans (*m/z* 350-1500) were acquired in the Orbitrap at 120,000 resolutions (at *m/z* = 400) in profile mode, with a maximum injection time of 100 msec and an AGC target of 400,000 ions. The S-lens RF level is set to 60. Isolation is performed in the quadrupole with a 1.6 Da isolation window, and HCD MS/MS acquisition is performed in profile mode using rapid scan rate with detection in the ion-trap, with the following settings: parent threshold = 5,000; collision energy = 32%; maximum injection time 56 msec; AGC target 500,000 ions. Monoisotopic precursor selection (MIPS) and charge state filtering were on, with charge states 2-6 included. Dynamic exclusion is used to remove selected precursor ions, with a +/- 10 ppm mass tolerance, for 15 sec after acquisition of one MS/MS spectrum.

### Database searching for mass spectrometry

Tandem mass spectra were extracted and charge state deconvoluted by Proteome Discoverer (Thermo Fisher, version 2.2.0388). Charge state deconvolution and deisotoping were not performed. All MS/MS samples were analyzed using Sequest (Thermo Fisher Scientific, San Jose, CA, USA; version IseNode in Proteome Discoverer 2.5.0.402). Sequest was set up to search a human and reformatted FASTA files assuming the digestion enzyme trypsin. Sequest was searched with a fragment ion mass tolerance of 0.60 Da and a parent ion tolerance of 10.0 PPM. Carbamidomethyl of cysteine was specified in Sequest as a fixed modification. Deamidated of asparagine and glutamine, oxidation of methionine and acetyl of the n-terminus were specified in Sequest as variable modifications

### Cell culture, and Transfection

Human neuroblastoma SH-SY5Y cells were purchased from ATCC (Cat # CRL-2266) and cultured in DMEM containing 15% FBS, 1% penicillin/streptomycin in CO2 incubator at 37°C. Plasmids encoding the human micro-RNA-adapted shRNA sequences targeting endogenous HTT, or BRG1 (Horizon Discovery, USA) were transfected into SH-SY5Y cells, and the transfected cells were selected for puromycin resistance. The knockdown efficiencies in the shRNA-expressing stable SH-SY5Y cells were assessed by western blots using appropriate antibodies. Plasmids expressing the full-length FLAG-tagged wild-type and mutant HTT cDNA (pRGS- wtHTT-Q19, pRGS-wtHTT-Q24, pRGS-mHTT-Q79, pRGS-mHTT-Q109 and pRGS-mHTT-Q145) were linearized with Pvu I and transfected into SH-SY5Y cells with Lipofectamine 2000 reagent (Invitrogen, USA), and the positive transfected cells were selected for G418 resistance, and the stable cells expressing the wtHTT or mHTT transgene were cultured in DMEM containing 15% FBS in CO2 incubator, and transgene expression was assessed by western blot with anti-FLAG or HTT antibodies. Mouse striatal cells STHdhQ7 and STHdhQ111^50^ were obtained from the Coriell Institute, USA, and cultured in DMEM containing 15% FBS, 1% penicillin-streptomycin, and in cell culture incubator at 33°C containing 5% CO2. The human PNKP and full-length HTT cDNA were PCR-amplified with appropriate primers, and sub-cloned into plasmid pSELECT-Puro-mcs (InvivoGen, USA) with appropriate linkers to construct pRGS- PNKP and pRGS-HTT-Q24, respectively. These plasmids were transfected into STHdhQ111 cells, and the stable clones were selected for puromycin resistance, and PNKP or HTT expressions in these cells were verified by western blots.

Plasmids encoding the human micro-RNA-adapted shRNA sequences targeting HTT, BRG1 or PNKP from the Horizon Discovery (USA) and were transfected into SH-SY5Y cells, and the transfected cells were selected for puromycin resistance. The knockdown efficiencies in the shRNA-expressing stable SH-SY5Y cells were assessed by western blots using appropriate antibodies. Cells were transfected with various plasmid DNA constructs using Lipofectamine 2000 reagent (Invitrogen, USA). Human embryonic kidney 293 (HEK293) cells were purchased from Coriell Institute, USA (Cat # CRL-1573) and cultured in EMEM, containing 10% FBS, 1% penicillin-streptomycin. HEK293 cells were co-transfected with plasmids p-FLAG-HTT and p- Myc-Ku70, p-Myc-Ku80, p-Myc-XRCC4, p-Myc-Ligase IV, p-Myc-PARP1 or Myc-ATM using Lipofectamine 2000 reagent (Invitrogen, USA). Plasmids expressing Myc-tagged BRG1 (p-Myc-BRG1), FLAG-tagged BRG1 (p-FLAG-BRG1) or the N-terminal truncated fragments (Exon 1) of HTT (p-FLAG-NT-HTT-[CAG]n carrying various lengths of polyQ sequences (Q15, Q19, Q40, Q42, Q79 or Q109) were transiently transfected into HEK293 cells and cells were cultured in MEM medium (Invitrogen, USA) containing 10% FBS and 1% penicillin-streptomycin. All the cell lines used in this study were authenticated by short tandem repeat analysis in the UTMB Molecular Genomics Core. We routinely tested mycoplasma contaminations in all our cell lines using GeM Mycoplasma Detection Kit (SIGMA, Cat # MP0025) and cells were found to be free from mycoplasma contamination.

### Antibodies and western blot analysis

Cell pellets or mouse brain tissues were homogenized, and total protein was isolated using a total protein extraction kit (Millipore, USA). The cytosolic and nuclear fractions were isolated from cells/tissue extract using a NE-PER nuclear protein extraction kit (Fisher Thermo Scientific, USA). WBs were performed according to the standard procedure, and each experiment was performed a minimum of three times to ensure reproducible and statistically significant results. The antibodies for APE1 (Cat # 4128), Ku80 (Cat # 2753), 53BP1 (Cat # 4937), phospho-53BP1 (Cat # 2675), p-ATM (Cat # 4526), rabbit polyclonal Ab for BRG1 (Cat # 3508), mouse monoclonal Ab for BRG1 (Cat # 52251), rabbit monoclonal Ab for ATM (Cat # 2873), and rabbit monoclonal Ab for DNA ligase IV (Cat # 14649) were from Cell Signaling Technology (USA); anti-p-ATM-S1981 rabbit monoclonal (Cat # ab81292) and anti-p-DNA-PKcs (Cat # ab18192) and monoclonal anti-PNKP (Cat # ab181107) were from Abcam, USA. Mouse monoclonal anti-HTT antibodies (Cat # MAB 2170 and Cat # 2166) were from Millipore-Sigma (USA). Rabbit monoclonal anti-HTT Ab (Cat # 5656) was from Cell Signaling Technology. Anti-Ku70 (Cat # SC-5309) XRCC4 (Cat # SC- 271087), CSB (Cat # SC-398022), ATM (Cat # SC-23921), BRG1 (Cat # SC-17796), PARP1 (SC-365315) and DNA-PKcs (Cat # SC-390849) mouse monoclonal antibodies were from Santa Cruz Biotechnology (USA). Anti-PNKP rabbit polyclonal antibody (Cat # MBP-1-A7257) and anti- BRG1 rabbit polyclonal antibody (Cat # NB100-2594) were from Novus Biologicals (USA), and the anti-p-PNKP (S114) rabbit polyclonal antibody (Cat # PA5-64846) was from Fisher Thermo Scientific (USA). Anti-FLAG M2 Ab (Cat # F3165) was from Sigma, and anti-Myc 9E10 Ab (Cat # SC-40; Santa Cruz) was from Santa Cruz Biotechnology (USA).

### Preparation of nuclear and cytosolic protein extracts

Whole-cell lysates and cytoplasmic and nuclear fractions were prepared as previously described. Mt extracts were prepared using an Mt extraction kit (Invitrogen, USA). Briefly, 5 × 10^6^ cells were ground with a glass homogenizer in an ice bath for 20 to 22 strokes. Cytoplasmic and nuclear fractions were separated by differential centrifugation (600 × g, 10 min, 4°C and 10,000 × g, 10 min, 4°C). The supernatant (cytosolic fraction) and pellet (mitochondrial fraction) were collected. The nuclear and cytosolic pellets were lysed to yield the final cytosolic and nuclear lysates, and the extracted proteins were analyzed by WBs.

### Analysis of HTT-associated TC-NHEJ proteins by co-immunoprecipitation (co-IP)

Co-IP analyses were performed using nuclear protein extracts of neuronal cells according to our reported protocol^19^. In brief, nuclei from neuronal cells were isolated using a nuclear protein isolation kit (Invitrogen, USA). Isolated nuclei were washed with phosphate-buffered saline (PBS), treated with trypsin (1 mg/ml in PBS) for 15 min at room temperature to remove contaminating proteins adhering to the outer nuclear surface, and washed extensively with ice-cold PBS. Nuclear protein extracts (NEs) were isolated using a nuclear protein IP kit (Sigma-Aldrich, USA) then treated with benzonase to remove DNA and RNA to avoid nucleic acid-mediated co-IP. Specific target proteins were immunoprecipitated and the resulting immunocomplexes (ICs) were washed extensively with Tris-buffered saline (TBS, 50 mM Tris-HCl [pH 7.5] and 200 mM NaCl) containing 1 mM EDTA, 1% Triton X-100, and 10% glycerol. The complexes were eluted from the beads with a solution of 25 mM Tris-HCl (pH 7.5), 500 mM NaCl, 2% SDS, then analyzed by WBs.

For co-IP from mouse brain tissue, approximately 250 mg cortex from freshly sacrificed WT mice was harvested; sliced into small pieces; collected in a sterile, pre-chilled, all-glass homogenizer; and mechanically homogenized with 4 volumes of ice-cold homogenization buffer (0.25 M sucrose, 15 mM Tris-HCl [pH 7.9], 60 mM KCl, 15 mM NaCl, 5 mM EDTA, 1 mM EGTA, 0.15 mM spermine, 0.5 mM spermidine, 1 mM dithiothreitol [DTT], 0.1 mM phenylmethylsulfonyl fluoride [PMSF] or protease inhibitors [Roche Applied Science, Germany]^19^. Homogenization was continued for ∼20 strokes or until a single-cell slurry was obtained (monitored under a microscope to ensure cell dissociation), incubated on ice for 15 min, and centrifuged at 1,000 × *g* to obtain the cell pellet. Nuclei were isolated as described above and the NEs were prepared for subsequent co-IP. Immunocomplexes (ICs) were analyzed by WBs for the presence of interacting protein partners using appropriate antibodies.

### Chromatin immunoprecipitation (ChIP)

The cells were grown on 10-cm tissue culture plates in DMEM containing 10% FBS. Formaldehyde was added to the medium to a final concentration of 1%, and the plates were incubated for 30 minutes. The cell pellets were collected and sonicated with a Fisher sonicator for twelve 15-s cycles at 60% output power; the lysates were centrifuged at 13,000 × *g* for 10 minutes and diluted 10-fold in ChIP dilution buffer. Genomic DNA sequences were detected with primers specific for the various genomic sequences e.g., Neurod1, Neurod2, NeuroG1. ChIP assays were performed using fresh mouse brain tissue as previously described^19^ with minor modifications. Briefly, 80-100 mg freshly harvested cortical tissues were chopped into small pieces (between 1 and 3 mm^3^) using a scalpel razor and fixed in 1% formaldehyde for 15 min with gentle agitation at room temperature to cross-link DNA to bound proteins. The samples were centrifuged at 440 × *g* for 5 min at room temperature, followed by the addition of 0.125 M glycine to terminate the cross-linking reaction. The samples were washed two or three times with ice-cold PBS (containing a protease inhibitor mixture) and centrifuged each time at 440 × *g* for 4 min at 4°C. The pellet was re-suspended in 1 ml ice-cold lysis buffer (10 mM EDTA, 1% [w/v] SDS, 50 mM Tris-HCl [pH 7.5]) with protease inhibitors and PMSF for 15 min on ice and homogenized slowly (∼20 strokes on ice) with a hand homogenizer and tight pestle to produce a single-cell suspension. After homogenization, the samples were transferred to pre-cooled 1.5-ml centrifuge tubes and centrifuged at 2260 × *g* for 5 min. The pellet was re-suspended in ice-cold lysis buffer (500 μl) and subjected to sonication to generate ∼500 bp of DNA fragments (sonicated 7× for 3 min for each pulse [21 min total]). The samples were centrifuged at 20,780 × *g* for 30 min at 4°C, and the supernatants were collected for ChIP as described previously (Gao et al. 2019). The sheared chromatin was immunoprecipitated for 6 h at 4°C with 10μg isotype control IgG (SC-2027; Santa Cruz Biotechnology, USA) or anti-HTT (MAB2170; Millipore-Sigma), antiKu70 (Cat # SC-5309; Santa Cruz Biotechnology, anti-XRCC4 (Cat # SC-271087; Santa Cruz Biotechnology), anti-DNA-PKcs (Cat # SC-390849; Santa Cruz Biotechnology), anti-DNA ligase IV (Cat # 52251; Cell Signaling Technology) or PNKP (Cat # MBP-1-A7257) antibodies. After DNA recovery with proteinase K treatment followed by phenol extraction and ethanol precipitation, 1% of input chromatin and the precipitated DNA were analyzed by qPCR with the following primers. ChIP data are presented as percent binding relative to the input value.

The following mouse and human-specific primer sets were used for the ChIP assays.

**Mouse Neurod1:** Neurogenic Differentiation Factor 1

F: CTGCAAAGGTTTGTCCCAAGC; R: CTGGTGCAGTCAGTTAGGGG

**Mouse Neurog1:** Neurogenin 1

F: GCTTGCTCCAGGAAGAACCT; R: AGAGACACCGCTACTAGGCA

**Mouse Tubb3:** Tubulin Beta 3 Class III

F: GTGGGGCTCTCCCCTAAAAC; R: TTGGGAGCGCACAGTTAGAG

**Human NEUROD1:** Neurogenic Differentiation Factor 1

F: GGTGCCTTGCTATTCTAAGACGC; R: GCAAAGCGTCTGAACGAAGGAG

**Human NEUROD2:** Neurogenic Differentiation Factor 2

F: GCTACTCCAAGACGCAGAAGCT; R: CACAGAGTCTGCACGTAGGACA

**Human BDNF:** Brain-derived neurotrophic factor

F: CATCCGAGGACAAGGTGGCTTG; R: GCCGAACTTTCTGGTCCTCATC

### Site specific DNA double strand break ChIP assay

As described previously, cells were synchronized transfected with I-SceI endonuclease expression vector or treated with the drug or liposomes as described^49^. Chromatin was centrifuged, the supernatant collected and diluted with ChIP dilution buffer as described previously. Diluted chromatin was incubated with specific antibody and the DNA was purified using QIAquick Spin columns (Qiagen). qPCR was carried out with specific sets of primers as shown in Table 1. Each experiment was repeated 3–4 times with consistent results. The signal to input ratio was low but significantly higher in comparison to IgG control values.

Chr1A (F) CTGAGTGTGGGAAGCTCTGG

Chr1A (R) GGAAGCAGTCTTCTCCGCTT

Chr1B (F) TACCCTCAGCTCCTGTACCC

Chr1B (R) GAGAGAAGCAAAGTTGCCAGCC

Chr17A (F) CTGCTCTCCAACAGGCCAG

Chr17A (R) CCCACCCCTCTTCTTTCCAG

Chr17B (F) AGAAGCAGTGACACGATGGA

Chr17B (R) GTGTCAAAGCAACTTGGGCC

The primers that were used for the ChIP assay for the mouse cells. Chromosolal location: Chromosome 15 - NC_000081.7

Forward: 5’-CGCTCTCTGAAGAACCTCCG

Reverse: 5’-GAAGGGCTTGTGTCCTCTCC

### *In situ* proximity ligation assays (PLAs)

Neuronal cells were plated on chamber slides and cultured in DMEM containing 10% FBS for 24 hours. Cells were briefly treated with DNA damaging agents either bleomycin (5µg/mL) or etoposide (15 µM) for 30 minutes and cells were immediately fixed with ice-cold 4% paraformaldehyde (PFA), permeabilized with 0.2% Tween-20, and washed with 1× PBS. The fixed cells were incubated with primary antibodies for HTT, BRG1 and various DSB repair proteins. Samples were subjected to PLAs using the Duolink PLA kit (O-Link Biosciences, Sweden). Nuclei were stained with DAPI (4’, 6-diamidino-2-phenylindole), and the PLA signals were visualized under a confocal microscope at 20× magnification.

### Chromosome aberrations

Analysis of G1, S, and G2 type chromosomal aberrations at metaphase. For G1-type chromosomal aberrations analysis, exponentially growing cells were exposed to 3 Gy of ionizing radiation, incubated for 18 h and metaphases collected after 3 h Colcemid treatment^88^. For S-phase-specific chromosome aberrations, exponentially growing cells (pulse-labeled with bromodeoxyuridine [BrdU] were irradiated with 2 Gy of IR and metaphases were harvested following irradiation and colcemid treatment. For G2-specific chromosomal aberrations, cells were irradiated with 1 Gy, and metaphases were collected after 3 h colcemid treatment ^88^. Chromosome aberrations scored at metaphase were examined before and after irradiation. One hundred metaphases were scored for each experiment and each experiment was repeated three to four times. Data were expressed as mean ± SD from three to four different experiments and were analyzed by two-tailed unpaired Student’s t test. (****p < 0.0001)

### Neutral comet assays

Neutral comet assays were performed using a Comet Assay Kit (Trevigen, USA). Cells were suspended in 85 μL ice-cold PBS and gently mixed with an equal volume of 1% low-melting-point agarose. The cell suspension was dropped onto an agarose layer and incubated in lysis buffer for 1 h. After lysis, slides were incubated in buffer containing 1 mM EDTA (pH 13) for 40 min and electrophoresed for 1 h. The slides were stained and analyzed with a fluorescence microscope.

### HD iPSC culture and differentiation

All the reagents were purchased from Fisher Scientific (Waltham, MA USA), unless otherwise specified. Three controls: CS25iCTR18n6, CS14iCTR20n6, CS83iCTR33n1 and three HD: CS87iHD50n7, CS03iHD53n3 and CS09iHD109n1 iPSC lines were derived and cultured as previously described on hESC-qualified Matrigel (HD iPSC Consortium, 2017 [28319609]). Once at 70% confluency, neural induction, and differentiation of neural progenitors with the addition of Activin A (Peprotech, USA), was performed as previously described in (Smith-Geater et al., 2020). Neuronal maturation was performed as previously described^51^ on Nunc™ 6 well plates. After 3 weeks of maturation, medium was removed and cells were washed once with PBS pH 7.4, without Mg^2+^ and Ca^2+^. Subsequently, cells were washed with 4°C PBS pH 7.4, without Mg^2+^ and Ca^2+^, scraped using a cell scraper, pipetted into a centrifuge tube, and centrifuged at 250 *x g* for 3 minutes. PBS was removed and samples were flash frozen in liquid nitrogen.

### CRISPR gRNA liposomes synthesis

Liposomes were prepared as described previously^49^. In brief liposome were dissolved in 3 mL ethanol. In order to label liposomes, fluorescent phospholipid lissamine rhodamine B 1,2- dihexadecanoyl-sn-glycero-3-phosphoethanolamine, triethylammonium salt (rhodamine-DHPE, Invitrogen) was incorporated in the membrane at 2% of the total lipid to all liposome formulations. Liposomes were sonicated until the desired size was achieved using Branson 1510 (Branson, Danbury, CT). Further CRISPR gRNA liposomes were prepared by adding 1 μg GeneArtTM PlatinumTM Cas9 nuclease (Thermo Scientific, Vilnius, Lithuania) to 400 ng gRNA in 50 μL serum free MEM medium. CRISPR gRNA liposomes’ size (164.4 nm, polydispersity index 0.15) and zeta potential (23.93 ± 1.48 mV) were assessed in triplicates by dynamic light scattering using Zetasizer instrument (Malvern, Worcestershire, UK). gRNA sequences for the chromosome loci are given below:

Chr1A: GCCCCCAAUAACAAAAUCGAguuuuagagcuagaaauagcaaguuaaaauaaggcuaguccguuaucaacuugaaaaaguggcaccgagucggugcuuuu

Chr17A:

GGAGGAUGGCCCAAUGUCGCguuuuagagcuagaaauagcaaguuaaaauaaggcuaguccguuaucaacuugaaaaaguggcaccgagucggugcuuuu

Chr17B:

GAGCUGAUCUCUCCUACCUUguuuuagagcuagaaauagcaaguuaaaauaaggcuaguccguuaucaacuugaaaaaguggcaccgagucggugcuuuu

### Non-homologous End-joining (NHEJ) and Homologous Recombination (HR) assay

The procedure for performing the DSB repair assay by NHEJ or HR is same as described previously^49^ We used two plasmid-based approach to measure NHEJ-mediated DSB repair proficiency, namely the standard I-SceI-based GFP reporter assay^49^ and a shuttle plasmid containing an unligatable DSB termini. The DSB repair assay by NHEJ or HR was performed by the previously described procedure ^49^. To perform the NHEJ assay, we used the EJ5 GFP-Puro plasmid DNA which was integrated at the different sites of chromosome 1 at Chr1A and Chr1B in H1299 cells. For the HR assay, the DR-GFP cassette was stably integrated at different chromosomal sites in H1299 cells as described previously^49^. Using flow cytometry, we measured the percentage of GFP-positive cells after inducing DSBs using the inducible I-SceI system at Chr1A and Chr1B^49^. In the NHEJ assay, the cells were transfected with either HTT-siRNA or control-siRNA, and the GFP fluorescence was measured in cells positive for GFP at chromosome 1 site B (Chr1B) and relative values were measured. In the HR assay, the cells positive for GFP are counted at sites (Chr1A, Chr1B) and plotted relative to the value obtained for Chr1A. The relative frequency of HR was determined by counting cells positive for GFP at the site of chromosome 1 A (Chr1A), which gave the maximum percentage of positive cells, and this is considered as 1.

### HD transgenic mice

The HD zQ175 transgenic mouse model expresses full-length mHTT from the endogenous mouse HTT promoter, and the mutant human HTT exon 1 carrying expanded CAG sequences was inserted into the mouse HTT locus by homologous recombination^32^. The heterozygous mice and control littermates (n = 4-5 pools of 2 animals per genotype) were sacrificed, and fresh brain tissues were used to isolate genomic DNA for proteins for WB analyses. For immunofluorescence assays, transgenic and control mice were deeply anesthetized and transcardially perfused with sterile PBS followed by 4% PFA in PBS. Brains were postfixed overnight in fixative solution and embedded in OCT. Slides with 4-μm-thick frozen sections were processed for immunostaining. Sacrifice and tissue collection were performed according to standard approved procedures following national guidelines and animal protocols.

### LA-QPCR analysis to assess DNA damage

Genomic DNAs were isolated from control untreated SH-SY5Y cells, SH-SY5Y cells expressing HTT-RNAi, HTT cDNA, BRG1 cDNA, or BRG1-RNAi using the DNeasy™ Blood & Tissue Kit (Qiagen, Cat # 69504). To minimize possible aerial oxidation during genomic DNA isolation, 2, 2, 6, 6-tetramethylpiperidine-*N*-oxyl was added to all solutions to a final concentration of 100 μM immediately before use. Similarly, genomic DNAs were isolated from STHdhQ7, STHdhQ111 or STHdhQ111 cells expressing either PNKP or HTT using identical procedure. Tissues were harvested from the cortex (CTX) of control and HD transgenic mice, and genomic DNA was extracted using the genomic-tip 20/G kit (Qiagen, Germany). LA-QPCR assays were carried out following an existing protocol^19, 89, 90^. Genomic DNA was quantified, and gene-specific LA-QPCR analyses were performed using Long Amp Taq DNA polymerase (NEB, USA). Various genomic loci were PCR-amplified from the actively transcribing genes in brain. The cycle numbers and DNA concentrations were standardized before each final reaction so that the reaction remained within the linear amplification range^19, 90^. The final PCR conditions were optimized at 94 °C for 30 s (94 °C for 30 s, 55–60 °C for 30 s depending on the oligo annealing temperature, 65 °C for 10 min) for 25 cycles and 65 °C for 10 min. Each reaction used 15 ng of DNA template, and the LA-QPCRs for all studied genes used the same stock of diluted DNA samples to avoid amplification variations due to sample preparation. A small DNA fragment for each gene was amplified to normalize large fragment amplification. The PCR conditions were 94 °C for 30 s, 54 °C for 20 s, 68 °C for 30 s for 25 cycles, and 68 °C for 5 min. Short PCR used 15 ng of the template from the same DNA aliquot. The amplified products were visualized on gels and quantified with the ImageJ software based on three independent replicate PCRs. The extent of damage was calculated according to our recently described method^19^. The following primers were used for the LA-QPCR to assess DNA damage in nuclear DNA in mouse brain tissues.

**Mouse GluN1:** Glutamate Ionotropic Receptor NMDA Type Subunit 1

Long: F: AACCAGTCCTCCTTCCCAGACTCAA; R: AAAGATTGAGGACCCACAGCCCA Short: F: AGGTGCTGAAGCGTATTGGGCGCTG; R: GCGCTCCAAGCATTTACGCCAAC

**Mouse GluR1:** Glutamate Ionotropic Receptor AMPA Type Subunit 1

Long: F: GGGTCTACCCACGACAACTAAAGCC; R: AATGCCCTCCTTGCCACACAGTTT Short: F: TGATGGGAGGGCTCGTGGGTAGG; R: GCCCCCGTGAGTTAGGCCAGATGCAATCA

**Mouse Rab3a:** RAS-Associated Protein

Long: F: CTGAGCCTGTAACTCTGCACCTGAC; R: AAGGTGCTAGTGGATCAAACGGTG Short: F: TCGTTTGCATAGAAAGCCCCGCCC; R: ACACCCGAGTTGCGCTCCCCTTGAT

**Mouse Tgoln2:** Trans-Golgi Network Protein 2

Long: F: ACCCTCCTCCATCCCCATGAGATAA; R: TGAGGACACAGGAAATGACTGCA Short: F: AGCAGCCGTTGTCTGTGTGTTCCG; R: AAGCAGCATCGGCAACGGAGAGGT

**Mouse Napg:** NSF Attachment Protein Gamma

Long: F: CGGTGGTGGAGCCCATTCAATTAGT; R: TCGTGAGGGAAGAGGGAGAAACCTT

Short: F: AGGTGAGGGACCTGTAGCTGATGCT; R: TGCCGTTGCTCTGCTCCGCTTCTC

**Mouse MyoD1:** Myogenic Differentiation 1

Long: F: ATAGACTTGACAGGCCCCGA; R: GGACCGTTTCACCTGCATTG Short: F: ATCTGACACTGGAGTCGCTTT; R: TTAGTCTCAGCTGCTGGTTCC

**Mouse MyoG;** Myogenin

Long: F: ACAAGCCTTTTCCGACCTGA; R: CCATGGCCAAGGCGACTTAT Short: F: GGCCACCAGAGCTAGAACAG; R: ATGAAGGCTGTGGACTTGGG

**Mouse Neurod1:** Neurogenic Differentiation Factor 1

Long: F: ATCAAGCACCACATAGGCAAACCAC; R: GCTCAGCATCAGCAACTCGGCT Short: F: GGGAGAGAGGCAAGCAGAAGAAGA; R: TAGGAAAGGCACCCATAGCCACT

**Mouse ENO2γ:** Enolase 2 gamma

Long: F: CTTGTTCTTCGGGGACCCTC; R: CATCCGTGTGCTTAAGGGGT Short: F: TAGGGGTGCCTAGTCCTGTC; R: GAGTGCTGGATGTGTGGTCA

**Mouse Bdnf:** Brain-derived neurotrophic factor

Long: F: ACCACTGTGCTGCATTCTTAGCACT; R; CCCAAGACCACTGCCATACAACTG Short: F: CCAGTTTTCTCCATGTGCTCAGGCT; R: TCTTTGTTGCTCACCTTTACGACAC

**Mouse Bcl2L11:** Bcl-2-Like Protein 11

Long: F: ATTCAGCCCCTGTCTCCTCCCTATC; R: ACCAGGCTGCTTCATTTCAGCTTTG Short: F: GTGGCATTCTTTGTCTTTGGGGCTG; R: CTTTGTAGAGGGATGCGGAGAGCA

The following primers were used for the LA-QPCR to assess damage in genomic DNA from SH- SY5Y cells expressing HTT-RNAi, BRG1-RNAi, control-RNAi, and control cells.

**Human ENO2γ:** Enolase 2 gamma

Long: F: ACGTGTGCTGCAAGCAATTT; R: CCTGAAACTCCCCTGACACC Short: F: GGTGAGCAATAAGCCAGCCT; R: CAGCTTGTTGCCAGCATGAG

**Human NEUROD1:** Neurogenic Differentiation Factor 1

Long: F: CCGCGCTTAGCATCACTAAC; R: TGGCACTGGTTCTGTGGTATT Short: F: TGCCTCTCCCTTGTTGAATGTAG; R: TTCTTTTTGGGGCCGCGTCT

**Human NEUROG1:** Neurogenin 1

Long: F: AACTCCAAGTACCCTCCAGGT; R: GTGATTTGACTGTGCACCCC Short: F: TTTCCCCCTCCCCTAGTGAG; R: GGGTCAGTTCTGAGCCAGTC

**Human BDNF:** Brain-derived neurotrophic factor

Long: F: TGTGAGGGAGGTCTACTTGGCAGAA; R: CCTTTCCCCCATCCTTGTGTTTCCA Short: F: TGAGATGTCAGAACATTTTCCCGTG; R: GGCTTTGAAGGGATTCTGTTGGGT

**Human BCL2L11:** Bcl-2-Like Protein 11

Long: F: GTTGCCTCTTTCTGGCCTTTGGTTG; R: ATGTCCTGACCCCAGGCTATACAGT Short: F: CATCCTTCTGCAAAGCTGGTCTCCA: R: AAGTCCCTGACAACATTCCCCTGC

The following primers were used for the LA-QPCR to assess DNA damage in genomic DNA isolated from *Drosophila* brain.

**Drosophila β-tubulin**

Long: F: GTATTCCTGCGCCAGGAGGATCG; R: CAGATGCTGGAGCTGCCTTTGGA Short: F: CGAGGGATACCTGTGAGCAGCTT; R: GTCACTTCTTGTGCTGCCATCGT

### Immunohistochemical analysis

The paraffin-embedded human brain sections were deparaffinized and analyzed by co-immunostaining with anti-HTT mouse monoclonal Ab (MAB2170; Millipore-Sigma), and anti- BRG1 rabbit monoclonal Ab (Cat # 52251; Cell Signaling Technology) to assess co-localization of HTT and BRG1. The human and mouse brain sections, and neuronal cells were analyzed by immunostaining with anti-p-53BP1 Antibody (4128; Cell Signaling) or γH2AX antibody (9718; Cell Signaling) for the detection of DSBs in nuclear genome. Nuclei were stained with DAPI (Molecular Probe, USA) and the sections were imaged under a confocal microscope.

### HD autopsy brain tissue samples

Human autopsy specimens (caudate nucleus) were obtained from the NIH NeuroBiobank, in accordance with local legislation and ethical rules. Control brain samples were collected from age-matched individuals without neurodegenerative disorders. The HD brain tissue samples were obtained from patients with HD who were clinically characterized based on the presence of chorea and motor, mood, and cognitive impairment. The molecular diagnosis of HD was established by analyzing genomic DNA extracted from peripheral blood using a combination of PCR and Southern blotting. The CAG repeat lengths in HTT were established by sequencing the expansion loci of the mutant allele. Human brain samples were dissected, and flash frozen in liquid nitrogen and stored at −80°C until further analysis. Genomic DNA isolated from the caudate of HD patients’ and age-matched control brain tissue (caudate). The HD patients’ brain tissues that are used for the DNA damage analyses are: (1) 56 years-old, male; grade 2; (2). 58 years-old, male; grade 2; (3) 53 years-old, male; grade 3; (4) 52 years-old, male; grade 3; (5) 85 years-old, male; grade 3; (6) 53 years-old, female; grade 3. Various genomic DNA segments PCR amplified, and the PCR products analyzed on agarose gels and the intensity of the DNA bands quantified using Image J. Five biological replicates and three technical replicates were used in this study.

### Image collection

The confocal images were collected using a Zeiss LSM-510 META confocal microscope with 40× or 60× 1.2 numerical aperture water immersion objectives. Images were obtained using two excitation wavelengths (488 and 543 nm) by sequential acquisition. Images were collected using 4-frame-Kallman-averaging with a pixel time of 1.26 μs, a pixel size of 110 nm, and optical slices of 1.0 μm. Z-stack acquisition was performed at 0.8-μm steps. Orthogonal views were processed with LSM 510 software.

## Statistical analysis

Data reported as mean ± SD and the statistical analysis was performed using Sigma Plot (SYSTAT Software). Differences between two experimental groups were analyzed by Student’s t test (2-tail, assuming unequal variances). When comparing multiple groups, One-way ANOVA was performed followed by Holm-Sidak test to determine significance. Statistical analyses of all data were performed using t test in GraphPad Prims Version 7.03 (*P < 0.005, ** P < 0.01, *** P < 0.001, **** P < 0.0001).

## HTT complex modeling

Due to the large size of HTT (∼350 kDa), alphafold models (HTT-BRG1, HTT-PNKP, HTT- KU70/80) were predicted separately using AlphaFold3 server (https://alphafoldserver.com/). Predicted local distance difference test (pLDDT) confidence score is shown in Figure 3G. The overall predicted template modeling (pTM) scores range between 0.61 and 0.69. A pTM score above 0.5 indicates that the overall predicted fold for the complex may be comparable to the true structure. Because polyQ expansion affects PNKP SUMOylation and activity based on^19^, the model with PNKP near the polyQ area was selected, results in the only non-clash HTT-BRG1- PNKP model out of 5 predicted models. Ku70/80 are also consistently predicted to bind on the N-HEAT domain of HTT with minimal clash with BRG1. PNKP-DNA model was generated by superimposed with murine PNKP-DNA crystal structure (PDB: 3zvn).

## Resource Availability

All data is available from the corresponding authors upon reasonable request.

## SUPPLEMENTARY FIGURE LEGENDS

**Supplementary figure 1.**
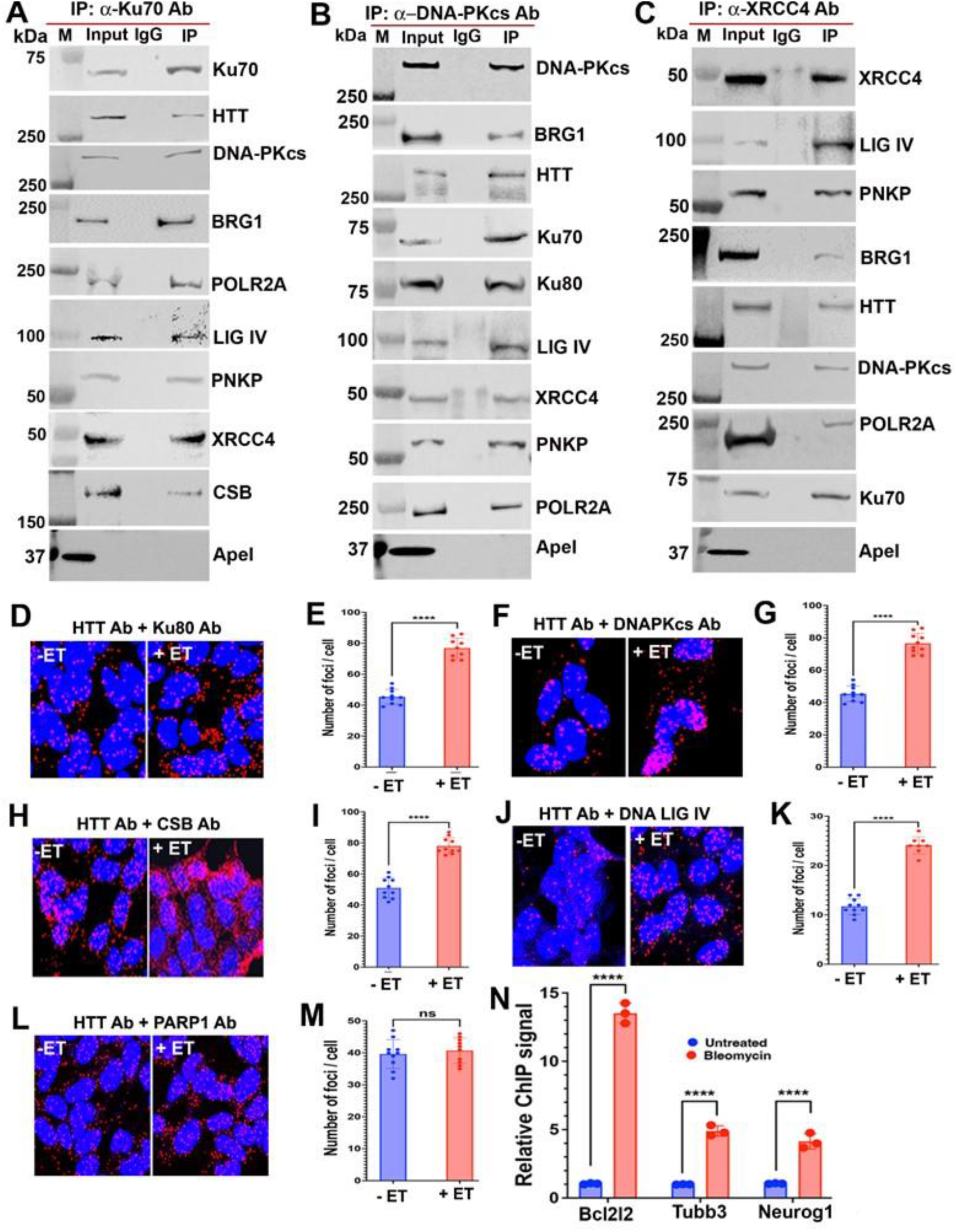
Characterization of the HTT-assembled transcription coupled non- homologous end-joining (TC-NHEJ) complex. **(A)** Nuclear extracts (NEs) isolated from 3-month-old C57BL/6 mouse brain, and Ku70 IP’d from the NEs with an anti-Ku70 mouse Ab (SC-5309; Santa Cruz), and the Ku70 immunocomplex (ICs) analyzed by western blots to detect the endogenous HTT, BRG1 and associated NHEJ components. An anti-ApeI Ab (4128; Cell Signaling) used as negative control in the WBs in panels A, B and C. Lane 1; protein molecular weight marker; lane 2: Input; lane 3: IgG control IP; lane 4: IP of Ku70 with anti-Ku70 Ab. **(B)** NEs isolated from 3-month-old C57BL/6 mouse brain, DNA-PKcs IP’d with an anti- DNA-PKcs Ab (SC-390849; Santa Cruz), and DNA-PKcs ICs analyzed by WBs to detect the presence of HTT, BRG1 and NHEJ complex components. Lane 1; protein molecular weight marker; lane 2: Input; lane 3: IgG control IP; lane 4: IP of DNA-PKcs with anti-DNA-PKcs Ab. **(C)** NEs isolated from 3-month-old C57BL/6 mouse brain, and XRCC4 IP’d from the NEs with anti-XRCC4 Ab (SC-271087; Santa Cruz), and XRCC4 ICs analyzed by WBs to detect the presence of HTT, BRG1 and associated NHEJ factors. Lane 1; protein molecular weight marker; lane 2: IP Input; lane 3: IgG control IP; lane 4: IP of XRCC4 with anti-XRCC4 Ab. Proximity ligation assay (PLA) performed to assess interaction of HTT and NHEJ proteins in SH-SY5Y cells, and in the SH-SY5Y cells after inducing DSBs by treating the cells with etoposide (ET; 15µM; 30 minutes). Generation of red fluorescence indicates representative positive protein-protein interactions. Nuclei stained with DAPI (panels 4). **(D)** PLA performed with anti-HTT mouse Ab (MAB2170; Sigma-Millipore) and anti- Ku80 rabbit Ab (2753; Cell Signaling) before and after treating SH-SY5Y cells with ET to assess interaction between HTT and Ku80 in untreated cells and in cells treated with ET (after inducing DSBs). **(E)** Relative PLA signals show interaction of HTT and Ku80 in control (-ET) cells and ET- treated cells. Data represents mean ± SE, ****p<0.0001. **(F)** PLA performed with anti-HTT rabbit Ab, (5656; Cell Signaling) and anti-DNA-PKcs mouse Ab (SC-390849; Santa Cruz) before and after treating the SH-SY5Y cells with ET to assess interaction between HTT and DNA-PKcs in response to increased DSBs. **(G)** Relative PLA signals show interaction of HTT and DNA-PKcs in control (-ET) cells and ET-treated cells. Data represents mean ± SE, ****p<0.0001. **(H)** PLA performed with anti-HTT rabbit Ab (5656; Cell Signaling) and anti-CSB mouse Ab (SC-398022; Santa Cruz) before and after treating SH-SY5Y cells with ET to assess interaction of HTT with CSB before and after inducing DSBs. **(I)** Relative PLA signals show interaction of HTT and CSB in control (-ET) cells and ET- treated cells. Data represents mean ± SE, ****p<0.0001. **(J)** PLA performed with anti-HTT mouse Ab (2760; Millipore-Sigma) and anti-DNA LIG IV (LIG IV) rabbit Ab (14649; Cell Signaling) before and after treating SH-SY5Y cells with ET to assess interaction of HTT with CSB before and after inducing DSBs. **(K)** Relative PLA signals show interaction of HTT and LIG IV in control untreated cells (- ET) and in ET-treated (+ET) cells. Data represents mean ± SE, ****p<0.0001. **(L)** PLA performed with anti-HTT rabbit Ab (5656; Cell Signaling) and anti-PARP1 mouse Ab (14649; Santa Cruz) before and after treating SH-SY5Y cells with ET to assess interaction of HTT with PARP1 before and after inducing DSBs. **(M)** Relative PLA signals showing interaction of HTT and PARP1 in control (-ET) cells and ET-treated cells. Data represents mean ± SE, ****p<0.0001. **(N)** SH-SY5Y cells treated with BL for 30 minutes, and ChIP performed to assess association of PNKP with genome in control and BL-treated cells. Relative ChIP values measured after normalization to control IgG. Data represents mean ± SE, ****p<0.0001.

**Supplementary figure 2.**
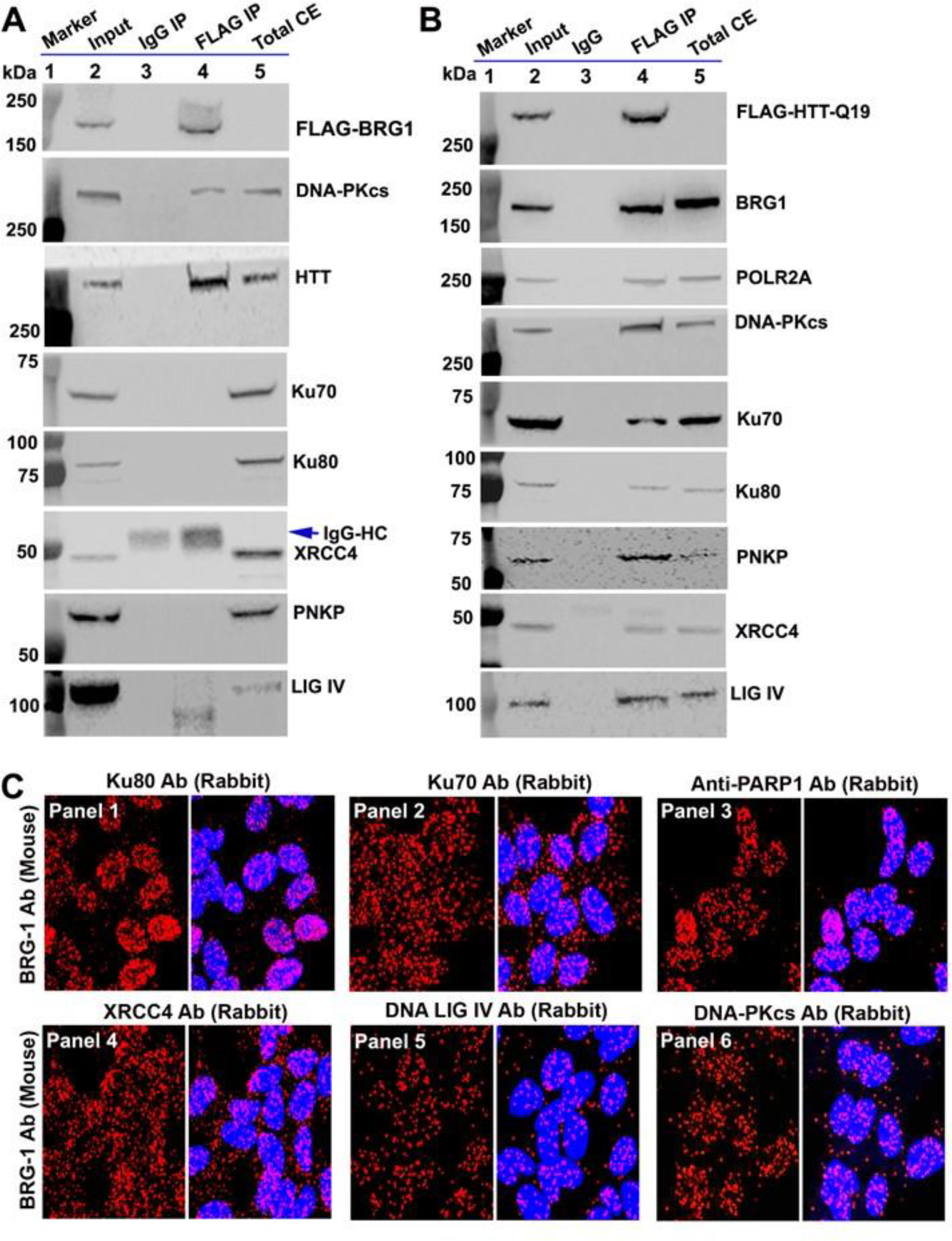
Characterization of the neuronal TC-NHEJ complex. **(A)** NEs isolated from SH-SY5Y cells constitutively expressing FLAG-tagged BRG1, and the exogenous FLAG-tagged BRG1 IP’d from the NEs with anti-FLAG Ab (F3165; Millipore-Sigma) and the FLAG IC analyzed by western blotting to detect HTT, and the key NHEJ proteins in the FLAG ICs. Lane 1: protein molecular weight marker; lane 2: Input; lane 3 IgG IP; lane 4: FLAG IP and lane 5: total cell extract (Total CE). The IgG heavy chain (IgG-HC) in the XRCC4 WB shown by arrow. **(B)** NEs isolated from SH-SY5Y cells constitutively expressing exogenous FLAG-tagged wtHTT carrying 19 glutamines (FLAG-wtHTT-Q19), and the exogenous FLAG- wtHTT-Q19 IP’d from the NEs with anti-FLAG Ab (F3165; Millipore-Sigma), and the FLAG ICs analyzed by western blotting to detect key NHEJ proteins in the FLAG ICs. Lane 1: protein molecular weight marker; lane 2: Input; lane 3 IgG IP; lane 4: FLAG IP and lane 5: total cell extract (Total CE). **(C)** Proximity ligation assay (PLA) performed to assess possible interaction of BRG1 and NHEJ proteins in SH-SY5Y cells. Generation of green fluorescence indicates representative positive protein-protein interactions. Nuclei stained with DAPI. PLA with anti-BRG1 (mouse) Ab and various NHEJ components e.g., Ku80 (Panel 1), Ku70 (Panel 2), PARP1 (Panel 3), XRCC4 (Panel 4), DNA ligase IV (Panel 5), DNA-PKcs (Panel 6), were performed before and after treating SH-SY5Y cells with ET.

**Supplementary figure 3.**
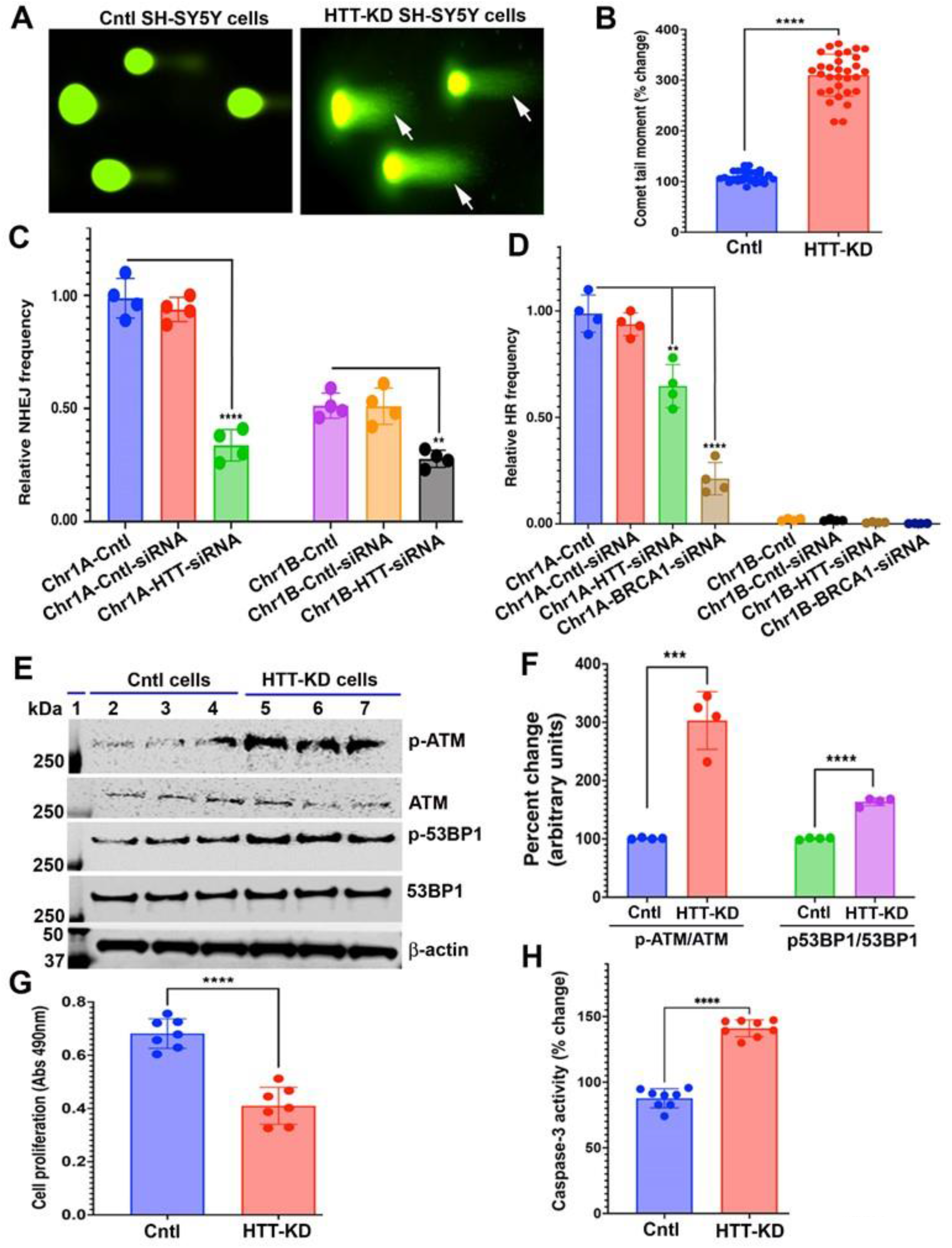
HTT depletion in neuronal cells impairs NHEJ-mediated DSB repair to induce DSBs. **(A)** Neutral comet analysis performed on control SH-SY5Y cells (left panel), and cells expressing HTT-RNAi (HTT-KD; right panel) to detect DSBs in genome. Comet tails indicate the presence of DSBs (arrows). **(B)** Relative DNA damage as analyzed by comet analysis in control (Cntl) and HTT- KD SH-SY5Y cells. Data represents mean ± SE. ****p<0.0001. **(C)** Depletion of HTT diminishes NHEJ frequency at both gene rich (Chr-1A) and gene poor (Chr-1B) sites. Data represents ± SD; **p<0.05; ****p<0.0001). **(D)** Homologous Recombination (HR) frequency at gene-rich Chr-1A site is reduced in HTT depleted (HTT-KD) cells. BRCA1 knockdown in SH-SY5Y cells used as positive control. Data represents mean ± SD; **p<0.05; ****p<0.001. **(E)** NEs isolated from the control SH-SY5Y cells (Cntl cells; lanes 2 to 4), and SH- SY5Y cells expressing HTT-RNAi (HTT-KD cells; lanes 5 to 7), and NEs analyzed by WBs to check the presence of DSBs by assessing activation of DNA damage- response pathway. WBs showing phosphorylation of ATM and 53BP1; β-actin used as loading control. Lane 1: Protein molecular weight marker in kDa. **(F)** Relative levels of the phosphorylation of ATM, and 53BP1 in control (Cntl) SH- SY5Y cells and in SH-SY5Y cells expressing HTT-RNAi (HTT-KD; HTT- knocked-down cells). Data represents mean ± SD, ***p<0.001; ****p<0.0001. **(G)** Cell viabilities (MTT assay) measured in control SH-SY5Y cells (Cntl), and SH- SY5Y cells expressing HTT-RNAi (HTT-KD cells). Data represents mean ± SD, ****p<0.0001. **(H)** Caspase-3 activities measured in SH-SY5Y cells expressing control-shRNA (Cntl) or HTT-shRNA (HTT-KD cells). Data represents mean ± SD, ****p<0.0001.

**Supplementary figure 4.**
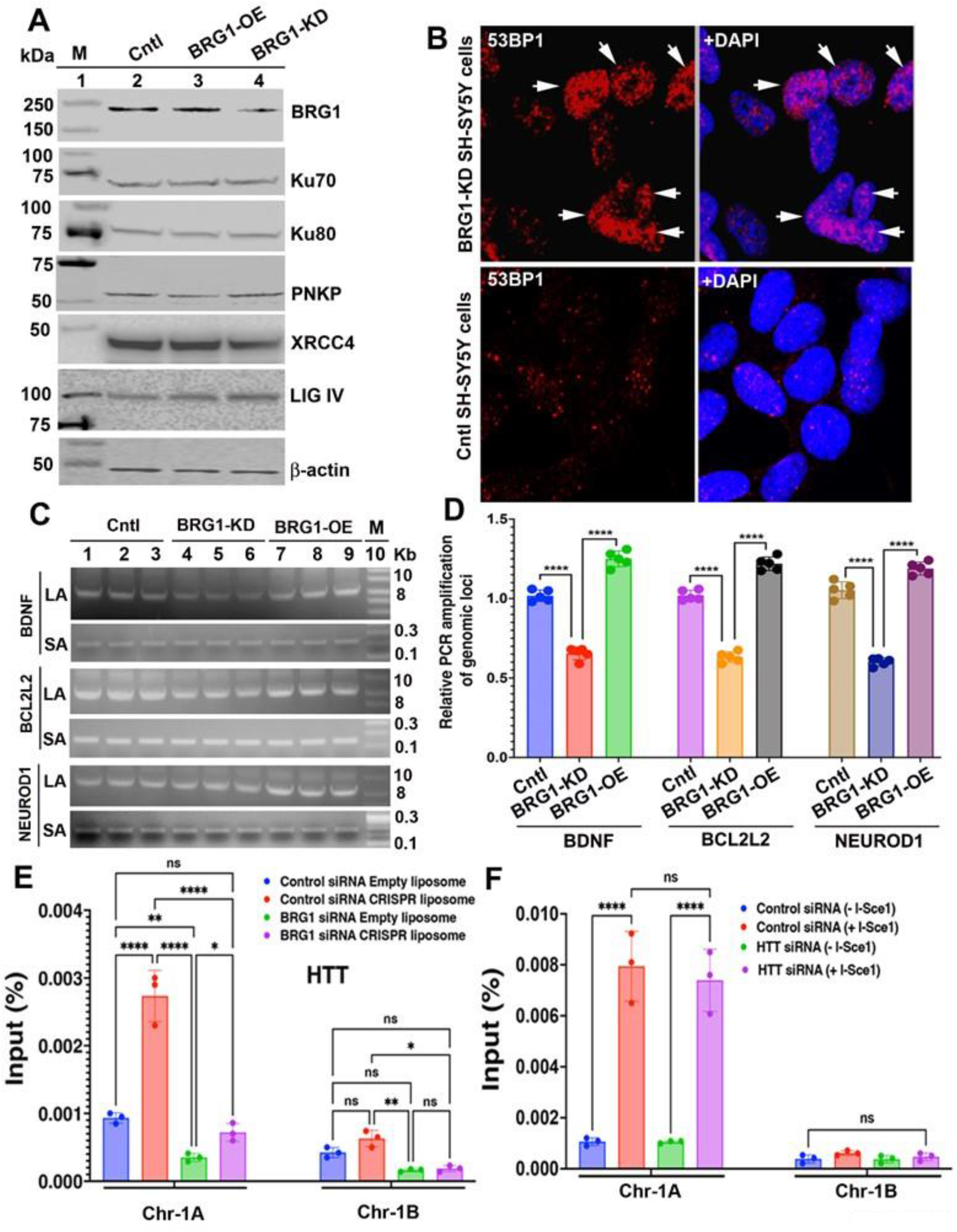
BRG1 regulates NHEJ-mediated DSB repair and maintains neuronal genome integrity. **(A)** NEs isolated from control SH-SY5Y cells (Cntl; lane 2), expressing human BRG1 cDNA (BRG1-OE; lane 3), expressing BRG1-RNAi (BRG1-KD; lane 4), NEs analyzed by WBs to detect BRG1, HTT and NHEJ protein levels; β-actin used as loading control. Lane 1: Protein molecular weight marker in kDa. **(B)** BRG1-KD SH-SY5Y cells (upper panel) analyzed by immunostaining with anti-53BP1 Ab (Cat # 2675; Cell Signaling) to detect the presence of DSBs (arrows). Nuclei stained with DAPI (blue). Control SH-SY5Y cells (lower panel) analyzed by immunostaining the cells with anti-53BP1 Ab (Cat # 2675; Cell Signaling) to detect the presence of DSBs in nuclei. Nuclei stained with DAPI (Blue). **(C)** Genomic DNAs isolated from the control SH-SY5Y cells (Cntl; lanes 1 to 3), SH- SY5Y cells expressing BRG1-RNAi (BRG1-KD; BRG1-knocked-down cells; lanes 4 to 6) or cells expressing human BRG1 cDNA (BRG1-OE cells; lanes 7 to 9), and DNA damage assessed by LA-QPCR. ∼8 to 10 kb regions of genome encompassing NEUROD1, BCL2L2, or BDNF) PCR-amplified, and the PCR products quantified. LA denotes long amplicon (8 to 10 kb); SA denotes short amplicon (0.2 to 0.3 kb); Lane 10: 1-kb DNA ladder. **(D)** Relative DNA damage in various genomic loci (NEUROD1, BCL2L2, or BDNF) quantified in control cells (Cntl), and cells expressing BRG1-RNAi (BRG1-KD) or human BRG1 cDNA (BRG1-OE). Data represents Mean ± SD. ****p<0.0001. **(E)** DSBs induced at transcriptionally active locus of chromosome 1 (Chr-1A) and transcriptionally inactive locus (Chr-1B) in control SH-SY5Y cells and BRG1-KD SH- SY5Y cells, cell harvested 135 minutes after adding liposomes, and ChIP performed to assess recruitment of HTT at the DSB sites in BRG1-KD and control SH-SY5Y cells. Data represents mean ± SD. ****p<0.0001; ns= not significant. **(F)** DSBs induced at transcriptionally active locus of chromosome 1 (Chr-1A) and transcriptionally inactive locus (Chr-1B) in control and HTT-KD SH-SY5Y cells, cell harvested 135 minutes after adding liposomes, and ChIP performed to assess the relative recruitment of BRG1 at the DSB site in HTT-KD and control SH-SY5Y cells. Data represents mean ± SD; ns= not significant.

**Supplementary figure 5.**
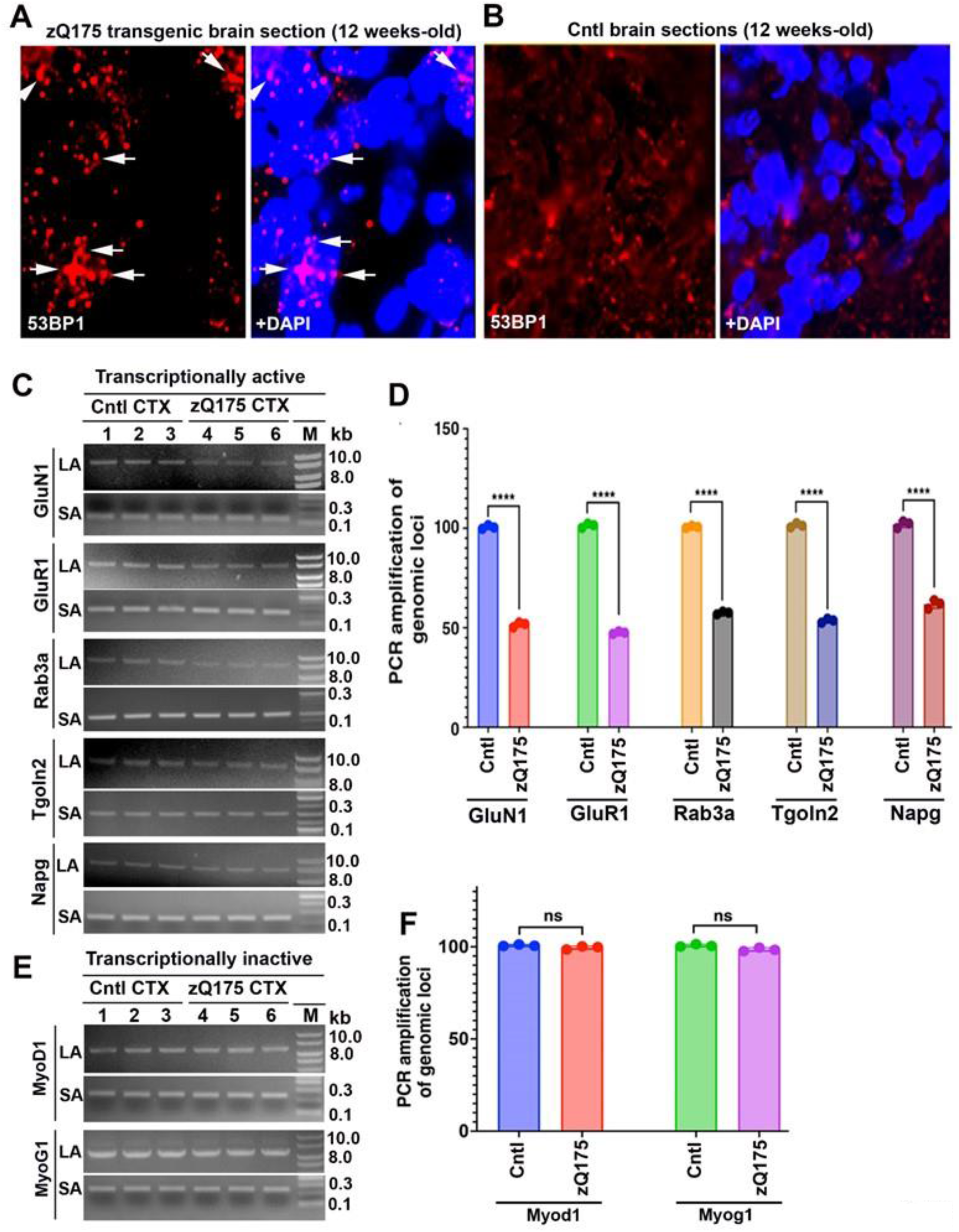
TC-NHEJ complex defects predominantly induce DSBs within the transcriptionally active genome. **(A)** Brain sections (cortex) from the zQ175 HD mice expressing mutant HTT (12-week-old heterozygous zQ175 mice) subjected to immunostaining with an anti-53BP1- S1778 Ab (4737; Cell Signaling) and analyzed by confocal microscopy. Nuclei stained with DAPI, and presence of DSBs in zQ175 brain sections shown by arrows. **(B)** Brain sections (cortex) from wildtype control mouse (12-week-old) subjected to immunostaining with an anti-53BP1-S1778 Ab (4937; Cell Signaling) and analyzed by confocal microscopy. Nuclei stained with DAPI. **(C)** Genomic DNAs isolated from the cortex (CTX) of 12-week-old zQ175 mice and control mice, and 6 to 8 kb regions from various transcriptionally active genomic DNA segments (GluN1, GluR1, Rab3a, Tgoln2 and Napg) PCR-amplified, and the PCR products from control mice (lanes 1 to 3) and zQ175 mice (lanes 4 to 6) analyzed, and intensity of the DNA bands quantified. LA: long amplicon (8 to 10-kb products), SA: short amplicon (0.2-0.3 kb products). Lane 7: 1 kb DNA ladder. **(D)** Relative PCR amplification of various transcribing gene loci (GluN1, GluR1, Rab3a, Tgoln2, Napg) in 12-week-old zQ175 and control mouse cortex (CTX). Data represent means ± SD, ****p<0.0001. Three biological replicates and three technical replicates were used for the analyses. **(E)** Relative PCR amplification of genomic DNA from the cortex (CTX) of pre- symptomatic zQ175 and control mice and genome segments that are transcriptionally inactive in brain (Myod1, and Myog1) PCR amplified and the PCR products from the control (lanes 1 to 3) and zQ175 mice (lanes 4 to 6) analyzed on agarose gels and quantified. Lane 7: DNA ladder in kb. **(F)** Relative PCR products from the transcriptionally inactive loci (Myod1 and Myog1) in the CTX of zQ175 transgenic and control mice. Three biological replicates and three technical replicates were used for the analyses. NS denotes not significant.

**Supplementary figure 6.**
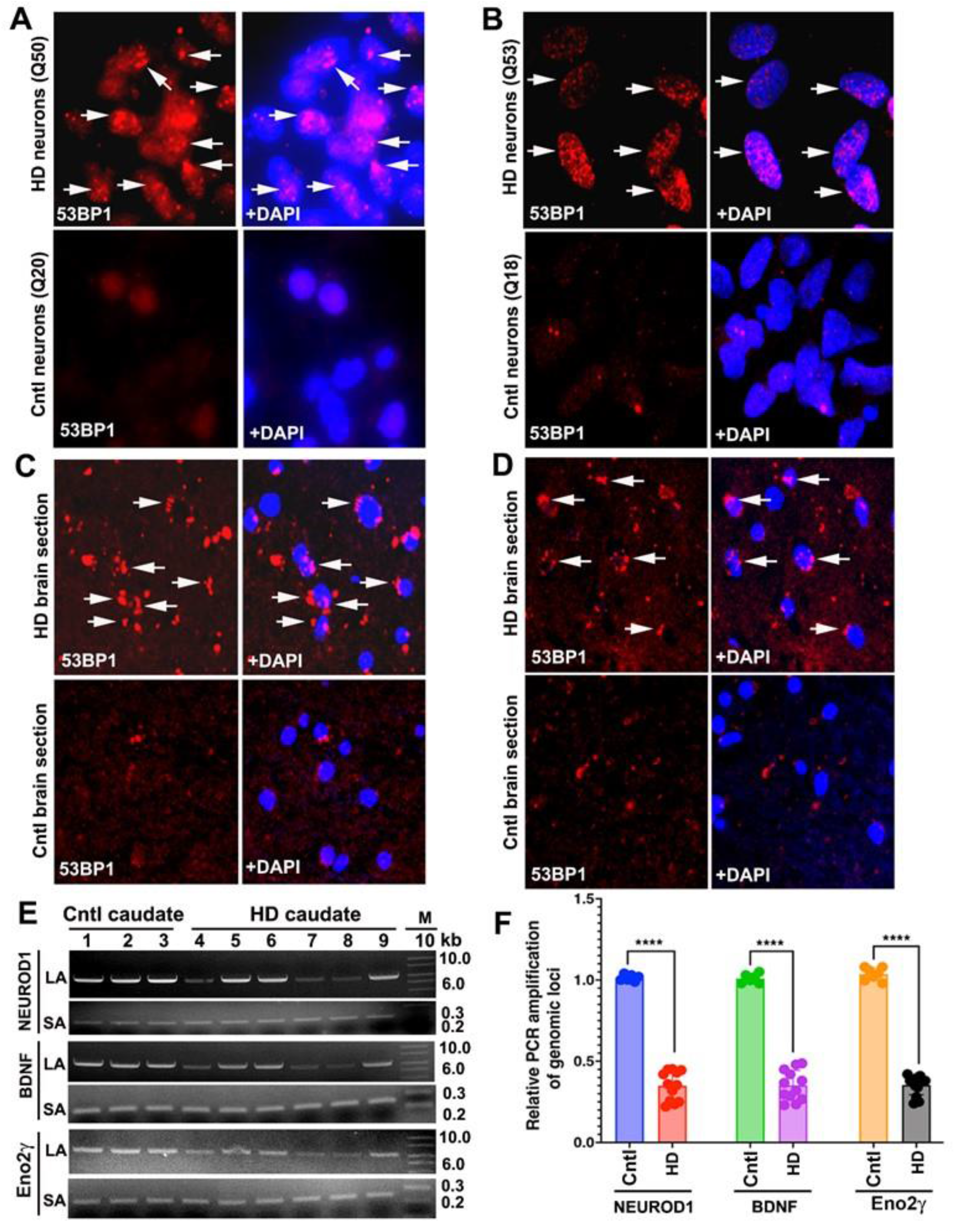
Impaired DNA repair results in persistence of DSBs in HD primary neurons and HD patients’ brain. **(A)** HD primary neurons (HTT encoding 50 glutamines: Q50; upper panel) and control neurons (encoding 20 glutamines: Q20; lower panel) analyzed to detect the presence of DSBs by immunostaining cells with an anti-53BP1 Ab (4937; Cell Signaling). Nuclei stained with DAPI, and the presence of DSBs in genomic DNA of mutant cells that appear as red puncta (upper panel) shown by arrows. **(B)** HD primary neurons (HTT encoding 53 glutamines: Q53; upper panel) and control neurons (HTT encoding 18 glutamines: Q18; lower panel) analyzed to detect the presence of DSBs by immunostaining the cells with an anti-53BP1 Ab (4937; Cell Signaling). Nuclei stained with DAPI, and presence of DSBs shown by arrows. **(C)** Brain sections (striatum) from HD patient (Age: 53 years; disease grade 2; upper panel) and age-matched control (lower panel; Age: 52 years) analyzed by immunostaining the sections with an anti-53BP1 Ab (4937; Cell Signaling), followed by confocal image analysis. Nuclei stained with DAPI (blue). The presence of 53BP1 Ab-positive DSBs in the HD brain nuclei (red fluorescence) is shown by arrows. **(D)** Brain sections (striatum) from HD patient (Age: 60 years; disease grade 3; upper panel) and age-matched control (lower panel; Age: 58 years) subjected to immunostaining with an anti-53BP1 Ab (4937; Cell Signaling) and followed by confocal image analysis. Nuclei stained with DAPI (blue). The presence of anti-53BP1-positive DSBs in the nuclei (red fluorescence) are shown by arrows. **(E)** Genomic DNAs isolated from HD patients’ caudate (lanes 4 to 9), and from age- matched control subjects (lanes 1 to 3) and DNA damage assessed using LA-QPCR. Genomic DNA segments encompassing NEUROD1, BDNF and ENOLASE from HD and controls subjects PCR-amplified and the PCR products quantified. LA denotes long amplicon; SA denotes short amplicon. Lane 10: 1-kb DNA ladder. HD patients’ brain tissues used for the DNA damage analyses: Lane 4: Age: 56-year-old, male; disease grade 2; lane 5: Age: 58-year-old, male; disease grade 2; lane 6: Age: 53-year-old, male; disease grade 3; lane 7: Age: 52-year-old, male; disease grade 3; lane 8: Age: 85- year-old, male; disease grade 3; and lane 9: Age: 53-year-old, female; disease grade 3. **(F)** Relative DNA damages in control (Cntl), and patients with HD. Three biological replicates for the controls and six biological replicates, and three technical replicates used for these calculations. Data represents mean ± SD; ****p<0.0001.

**Supplementary figure 7.**
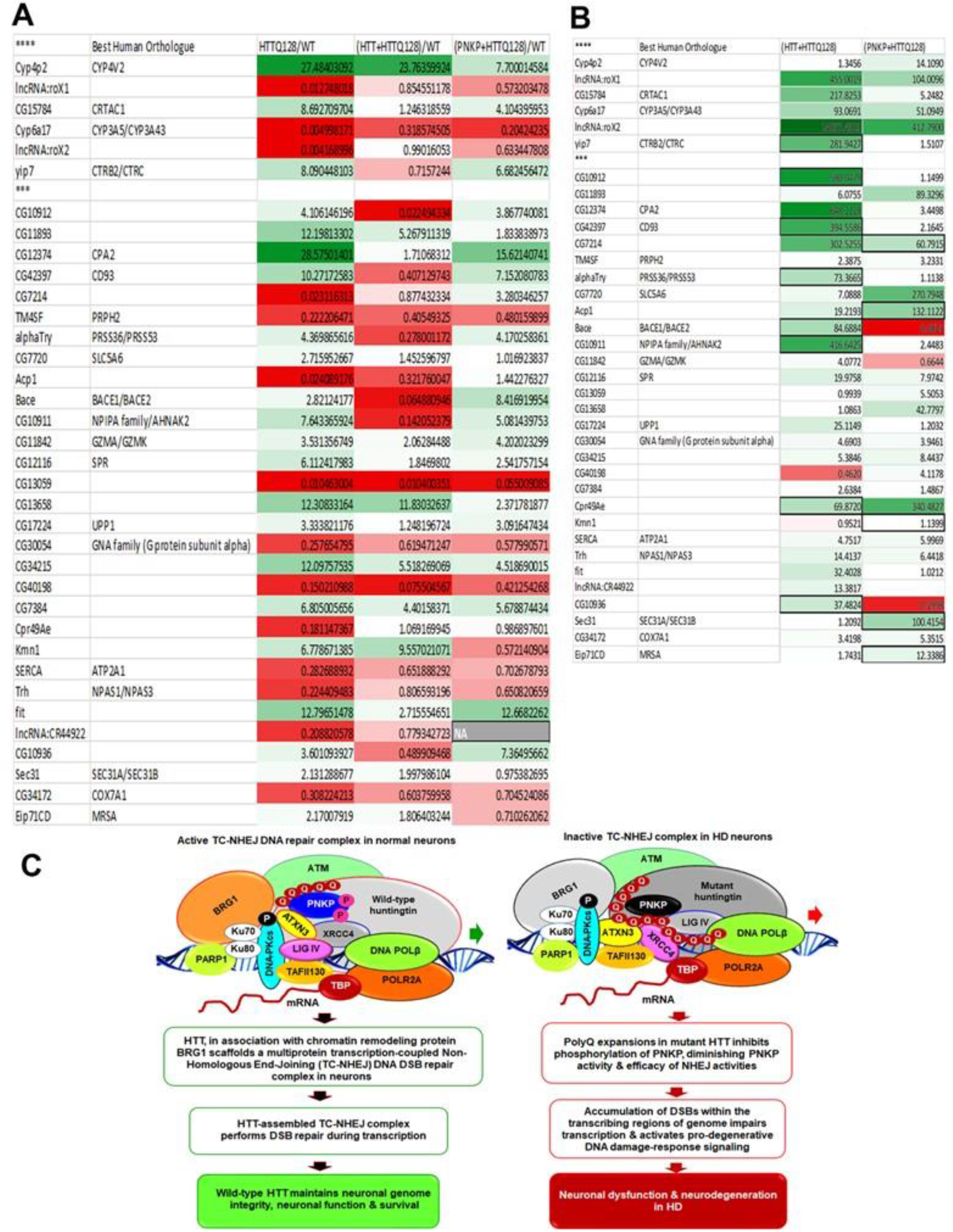
Ectopic expression of wtHTT or PNKP in Drosophila model of HD reverses the transcriptional landscape. **(A)** Table of genes identified in RNAseq analysis that have P values of less than 0.001 when *Drosophila* expressing mHTT (UAS-128Qhtt^FL^) were compared to WT control. The table depicts the genes sorted by increasing P value of difference when *Drosophila* expressing mHTT (UAS-128Qhtt^FL^) were compared to WT control with P < 0.0001 noted by **** and P < 0.0001 noted by ***. The color of each table cell is based on the variance value of detected RNA level for that gene (row) and mutant (column) compared to WT control with white as no change, green as increased levels, and red as decreased levels. The deeper the color the more the variance from the WT control levels. The variance value is also included in each cell. **(B)** The table depicts the weighted difference of the level of the co-overexpression compared to mHTT only expression. The color of each table cell is based on the weighted variance value of difference to WT level for that gene (row) and mutant (column) compared to mHTT only expression, with white as similar difference to WT control levels, green as more like WT control levels, and red as less as the WT control levels. The deeper the color the greater the difference in variance to mHTT only expression compared to the WT control levels. The weighted variance value is also included in each cell, and again higher value means closer to WT mRNA level. If the expression level compared to WT level flipped from greater than WT level to less than WT level or vice versa compared to mHTT expression the cell border is bolded. **(C)** Proposed schematic mechanisms whereby wild-type HTT (wtHTT) stimulates DNA repair to maintain genome integrity and how mutant HTT (mHTT) synchronously disrupts DNA repair and transcription in HD to trigger early neurotoxicity in HD. wtHTT and BRG1 assemble a DSB repair complex with essential NHEJ factors including PNKP, Ku70, Ku80, DNA-PKcs, XRCC4, DNA ligase IV, CSB and PNKP in neurons. This structure senses DSBs during transcription and orchestrates their repair. wtHTT thus plays a pivotal role in DNA repair and in maintaining genome integrity during transcription. In contrast, HTT polyQ expansions inhibit recruitment of various NHEJ factors at the DSB sites, degrading normal TC- NHEJ function and DSB repair. This leads to persistence of DSBs in genome and chronic activation of the DNA-damage-response in HD. Cumulative accumulation of DSBs within transcriptionally active genome adversely impacts expression of neuronal genes, amplifying pro-degenerative impacts. Mutant HTT thus synchronously impairs DNA repair and transcription, triggering neurotoxicity and functional decline in HD.

## Notes

### Competing Interest Statement

The authors have declared no competing interest.

